# scMODAL: A general deep learning framework for comprehensive single-cell multi-omics data alignment with feature links

**DOI:** 10.1101/2024.10.01.616142

**Authors:** Gefei Wang, Jia Zhao, Yingxin Lin, Tianyu Liu, Yize Zhao, Hongyu Zhao

**Affiliations:** Department of Biostatistics, Yale University, CT, USA; Program of Computational Biology and Bioinformatics, Yale University, CT, USA

## Abstract

Recent advancements in single-cell technologies have enabled comprehensive characterization of cellular states through transcriptomic, epigenomic, and proteomic profiling at single-cell resolution. These technologies have significantly deepened our understanding of cell functions and disease mechanisms from various omics perspectives. As these technologies evolve rapidly and data resources expand, there is a growing need for computational methods that can integrate information from different modalities to facilitate joint analysis of single-cell multi-omics data. However, integrating single-cell omics datasets presents unique challenges due to varied feature correlations and technology-specific limitations. To address these challenges, we introduce scMODAL, a deep learning framework tailored for single-cell multi-omics data alignment using feature links. scMODAL integrates datasets with limited known positively correlated features, leveraging neural networks and generative adversarial networks to align cell embeddings and preserve feature topology. Our experiments demonstrate scMODAL’s effectiveness in removing unwanted variation, preserving biological information, and accurately identifying cell subpopulations across diverse datasets. scMODAL not only advances integration tasks but also supports downstream analyses such as feature imputation and feature relationship inference, offering a robust solution for advancing single-cell multi-omics research.

## Introduction

Recent advances in single-cell technologies, which enable the measurement of transcriptomic [1], epigenomic [2, 3] and proteomic [4, 5, 6] profiles at single-cell resolution, have greatly enhanced our ability to comprehensively characterize cellular states. Data resources generated by these technologies have provided significant insights into the functions of various cell types [7, 8] and deeper understanding of pathology [9, 10] from multiple omics perspectives. As high-throughput single-cell technologies continue to develop rapidly and data resources accumulate, there is an increasing need for computational methods that can integrate information from different modalities to perform joint analysis of single-cell multi-omics data and gain a more comprehensive understanding of cellular states and functions.

However, integrating single-cell omics datasets presents unique challenges. First, cross-modality integration, also known as “diagonal integration” [11], aims to align different single-cell modalities with distinct features. For the integration of single-cell RNA-sequencing (scRNA-seq) and single-cell sequencing assay for transposase-accessible chromatin (scATAC-seq) datasets, the cross-modality features exhibit strong connections as gene expression levels can usually be accurately imputed using single-cell chromatin accessibility [12, 13]. Nevertheless, the features across some modalities, such as surface protein abundance in proteomic assays and its coding gene expression in scRNA-seq data, show weaker relationships which are often not robust enough to reliably guide integration, as mRNA levels do not always correlate with protein abundance due to post-transcriptional regulation, degradation, and protein modifications [14, 15, 16, 17]. Furthermore, many cross-omics features are involved in regulatory circuits that are not well understood, making it difficult to achieve integration when known information about feature relationships is limited. Second, compared to scRNA-seq which provides whole-transcriptome profiling for tens of thousands of genes, some technologies detect only a limited number of features, such as dozens to hundreds of protein targets in antibody-based single-cell proteomics [5, 18] and 100 to 1,000 genes in imaging-based spatial transcriptomics [19]. This limitation further constrains the signal available for high-quality integration, making cross-modality integration more challenging.

Many computational methods have been developed for the integration of single-cell datasets [20, 21, 22, 23, 24, 25, 26]. However, most of these methods were developed primarily for correcting batch effects in scRNA-seq datasets, or integrating omics layers with strong connections such as scRNA-seq and scATAC-seq data. These methods, however, often fail to address the aforementioned challenges. Among the existing methods, bindSC [25] and MaxFuse [26] were recently developed for single-cell multi-modal integration, demonstrating particular efficacy in integrating modalities with weak relationships, such as protein abundances and gene expression levels. Both methods utilize canonical correlation analysis (CCA) to learn linear projections that map features from each modality to a common space, ensuring that the projected vectors are maximally correlated. However, the inherent structure of unwanted variation across single-cell datasets is often complex and nonlinear [27, 28, 29]. Meanwhile, the relationships between cross-modality features can be intricate and cell type-specific, regulated by multiple biological factors [15, 16]. Thus, linear projections may lack the flexibility needed to adequately correct unwanted variation and accurately model feature correspondence.

Here, we present scMODAL, a general deep learning framework for single-cell multi-omics data alignment with feature links. scMODAL is designed to integrate unpaired datasets with limited numbers of known positively correlated features, which are also referred as “linked” features in the literature [26]. To capture complex relationship between different modalities, we build neural networks to project different single-cell datasets into a common low-dimensional latent space and apply generative adversarial networks (GANs) [30] to align cell embeddings. To accurately find cell population correspondence across datasets, scMODAL utilizes prior information from known linked features to identify anchor cell pairs that can guide integration, while preserving topology structure of all input features. Through comprehensive real data experiments, we demonstrate scMODAL’s performance in preserving biological variation across modalities and finding correct correspondences among them, using scRNA-seq, single-cell proteomics and scATAC-seq datasets. Especially, scMODAL shows state-of-the-art performance in both unwanted variation removal and biological information preservation even when there are very few linked features. With the integration results, scMODAL can identify cell subpopulations that were not distinguishable with the original modality features. We further showcase scMODAL’s capabilities in downstream tasks, such as imputation of cross-modality features and inference of feature relationships. We have made scMODAL publicly available as a Python package at https://github.com/gefeiwang/scMODAL.

## Results

### Method overview

scMODAL is a deep generative framework that learns integrated cell representations from single-cell multi-omics features. The input to scMODAL comprises cell-by-feature data matrices. For simplicity, we consider the scenario involving two datasets with different numbers of cells and features, denoted by 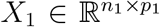 and 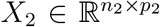. Using prior knowledge about the cross-modality feature relationships, we compile linked features from these datasets into another pair of matrices 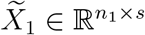 and 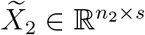, where *s* represents the number of feature pairs (Fig. 1**a**). The columns of these matrices pair cell features likely to be positively related, such as gene expression levels from scRNA-seq data and gene activity scores computed based on scATAC-seq data, or protein abundance levels paired with their corresponding protein-coding gene expression levels.

**Figure 1.**
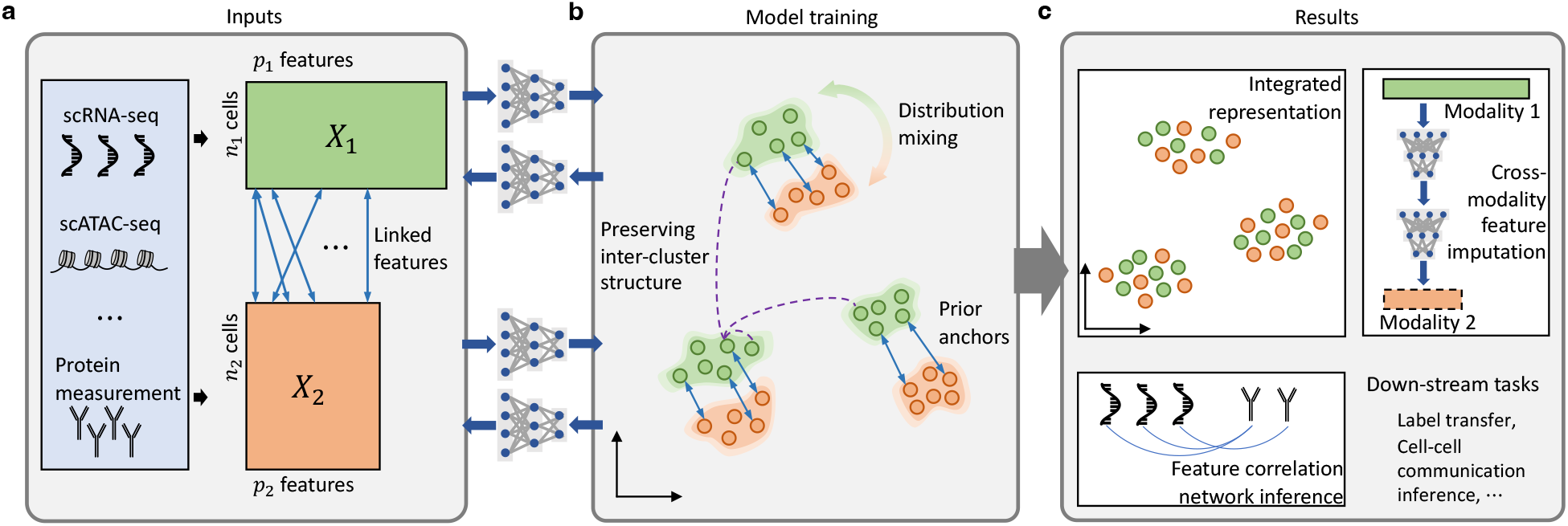
Overview of scMODAL. **a**. scMODAL takes single-cell feature matrices from different modalities, together with feature links as input. **b**. scMODAL utilizes generative adversarial learning to mix the distributions of cell embeddings from different datasets. To find correct correspondence between modalities as well as preserve biological variation within each modality, regularizations to narrow the distance between anchors based on prior information and preserve geometric representation of cells are applied in the training process of scMODAL. **c**. scMODAL outputs integrated cell representations for further analyses, and the composition of trained networks enables imputation of features and inference of feature relationship across single-cell modalities. The results can also be used for multiple downstream analyses, including label transfer for revealing cell identities and cell-cell communication inference using imputed features.

To address complex unwanted variations between modalities, we use nonlinear neural networks as encoders, denoted as *E*_1_ and *E*_2_, to map cells to a shared latent space *Z* (Fig. 1**b**). Unlike most integration methods that rely solely on shared features, our approach inputs the full feature matrices *X*_1_ and *X*_2_ into the encoders to preserve biological information. Decoders *G*_1_ and *G*_2_ are employed to generate cell features from the latent embeddings and trained together with the encoders for autoencoding consistency. Once the cells are encoded in *Z*, we apply the generative adversarial learning mechanism in GANs to minimize the Jensen-Shannon divergence between the latent distributions of the datasets using an auxiliary discriminator network.

However, using generative adversarial learning to align distributions without guidance can result in incorrect integration by mismatching distinct cell populations. In practice, there are often no cells measured with both modalities available to serve as integration anchors. Therefore, we use cell similarity information in positively related features 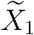 and 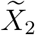 to establish connections between datasets. Specifically, during training, we calculate mutual nearest neighborhood (MNN) pairs between minibatches of samples as anchors to guide integration. After identifying these MNN pairs, we regularize the neural network optimization by keeping the embeddings of MNN pairs close to each other using an L2 penalty on the Euclidean distance. While using MNN pairs for batch-effect correction in scRNA-seq datasets has yielded promising results [27], simply minimizing the distances between MNN pairs may not effectively align all cell populations in a multi-omics setting, as the shared information between cross-modality features could be limited. Nevertheless, these MNN pairs can serve as valuable prior information, enhancing the accuracy of integration when combined with the generative adversarial learning mechanism. Additionally, to prevent the networks from becoming too flexible, which could result in loss of information and destruction of dataset-unique structures, we preserve the geometric structure of each dataset by regularizing the geometric representations of cells. Specifically, for each cell, we calculate its Gaussian kernel distance from other cells in the sampled minibatch as a *B*-dimensional geometric representation, where *B* is the batch size. During training, the encoders are encouraged to preserve the geometric representations, maintaining relative similarities and distinctions among cell populations.

After training the neural networks, aligned cell representations can facilitate cross-modality integrative analysis (Fig. 1**c**). The network compositions *E*_1_(*G*_2_(*·*)) and *E*_2_(*G*_1_(*·*)) can be used to map cells from one modality to another, serving as a bridge for cross-modality feature imputation. Using imputed features, we can also infer correlation networks among different modalities to reveal potential regulatory relationships. More details are provided in the Methods section.

### Benchmarking on integration of gene expression and protein abundance with CITE-seq PBMC data

We first evaluated scMODAL’s performance using a human cellular indexing of transcriptomes and epitopes by sequencing (CITE-seq) peripheral blood mononuclear cells (PBMCs) dataset [31], which simultaneously quantified transcriptome-wide gene expressions and 228 surface protein markers using antibody-derived tags (ADTs) in the same cells. We applied scMODAL and other recently developed integration methods, including MaxFuse [26], bindSC [25], GLUE [24], Portal [23] and Seurat [20], to integrate the RNA and ADT modalities, treating these cells as unmatched during the integration process. The matched RNA and ADT profiles in this dataset serve as the ground truth for a systematic comparison.

Before integration, we investigated the cell population structures in unintegrated datasets. As shown in the UMAP [32] plot and correlation heatmap based on the ADT data, CD4 T cells and CD8 T cells exhibit distinct protein abundance levels, indicated by their separate clusters and distinct correlation blocks (Fig. 2**a, c**). However, they show higher similarity when comparing the expression levels of highly variable genes (Fig. 2**d**).

**Figure 2.**
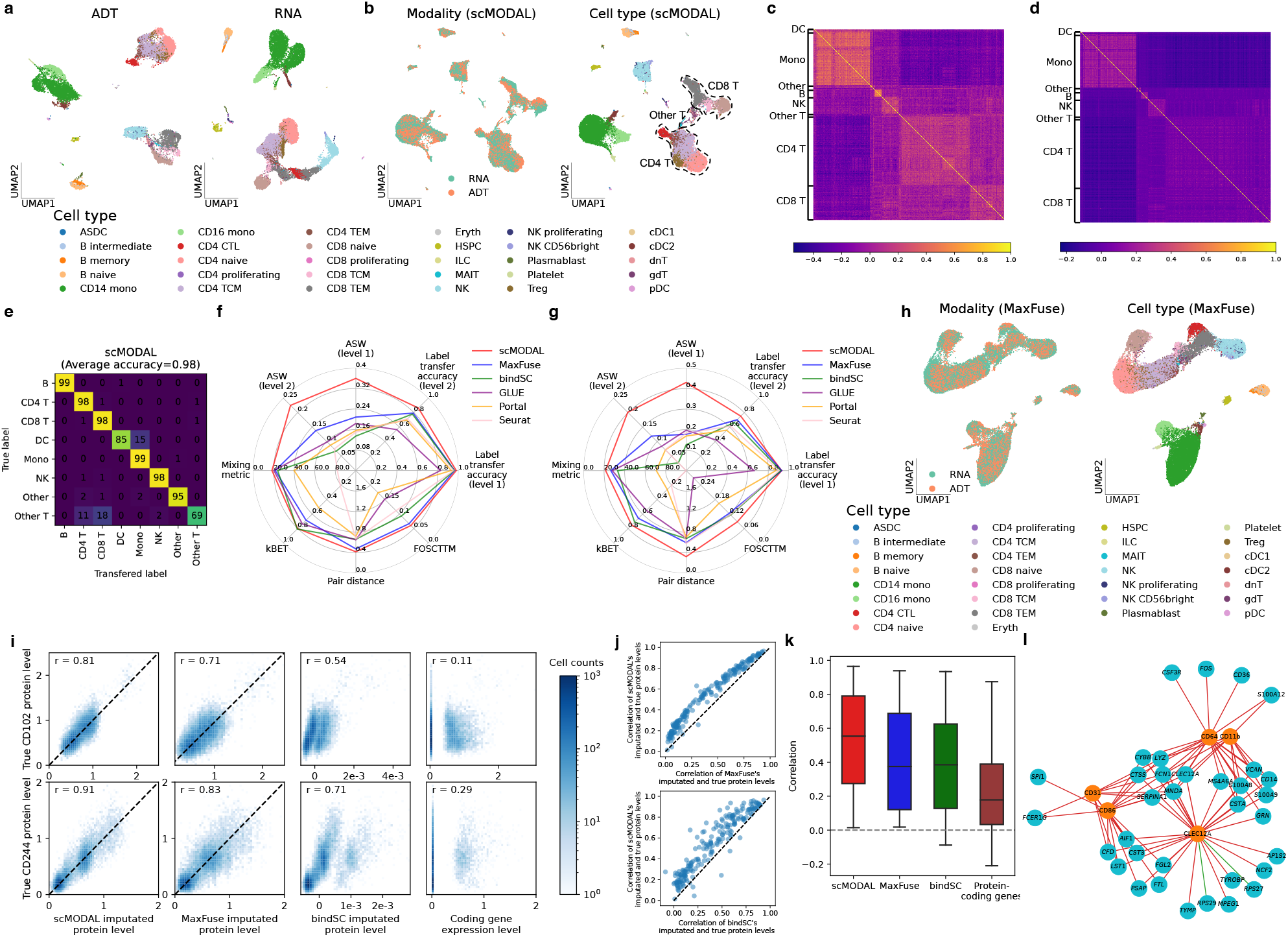
Benchmarking on integrating transcriptome and protein data produced by CITE-seq. **a**. UMAP plots of unintegrated ADT and RNA datasets, colored by cell types. **b**. UMAP plots of integrated embeddings produced by scMODAL, colored by modalities (left) and cell types (right). **c, d**. Correlation heatmaps of ADT (**c**) and RNA features (**d**). **e**. Confusion matrix of scMODAL’s label transfer result. **f, g**. Quantitative evaluations in terms of unwanted variation removal, biological information preservation and cell-state matching accuracy, using full protein panel (**f**) and 30 proteins (**g**). **h**. UMAP plots of integrated embeddings produced by MaxFuse, which ranked the second in preserving cell population structures, colored by modalities (left) and cell types (right). **i**. Comparison of scMODAL, MaxFuse and bindSC’s imputation results and coding genes of proteins CD102 and CD244. **j**. Comparison of scMODAL, MaxFuse and bindSC’s protein imputation results using Pearson correlation coefficient. **k**. Correlation box plots for scMODAL, MaxFuse and bindSC’s imputation results and protein coding genes, with the central lines marking the median values. **l**. Gene-protein correlation network inferred using scMODAL’s imputation result, centered at five monocyte-enriched proteins.

Among all compared methods, scMODAL, MaxFuse, bindSC and GLUE were specifically developed for integrating different single-cell modalities and can utilize dataset-unique features for integration, while Seurat and Portal only use linked features between modalities. We assessed the integration performance of these methods from three main aspects. First, a good integration method should mix the cell distributions well in its output. We used the mixing metric [20] and *k*-nearest-neighbor batch-effect test (kBET) [33] scores to indicate how the datasets are mixed after integration. Second, distinct cell types should be kept separated after integration. Using two levels of cell-type annotations, we quantified how different cell states are prevented from being incorrectly mixed together with the average silhouette width (ASW). Third, we measured how accurately corresponding cell states are matched between modalities using label transfer accuracy with labels transferred from RNA cells to ADT cells, relative distance between ground truth paired cells (pair distance), and fraction of samples closer than true match (FOSCTTM) [34]. More details about the metrics are provided in the Methods section.

We first inspected the ability of integration to mix cell distributions. As shown by the results (Fig. 2**f** and Supplementary Fig. 1), scMODAL achieved comparable alignment performance with cross-modality integration methods including MaxFuse, bindSC and GLUE, indicating its ability of removing strong cross-modality unwanted variation. Among compared methods, scMODAL has the best performance in integration accuracy. Notably, scMODAL had the highest label transfer accuracy scores among all methods, approximately 98% for level 1 annotation and 86% for level 2 annotation (Fig. 2**e, f**). Higher label transfer accuracy scores indicate that scMODAL is better at finding correct correspondence between cell states across RNA and ADT modalities. Meanwhile, scMODAL’s lower pair distance and FOSCTTM scores indicate that ground truth cell pairs have closer relative distances in its integrated embeddings compared to other methods. More importantly, scMODAL achieved significantly improved ASW scores compared to other methods, indicating its capability to preserve fine-grained cell populations. This result is consistent with our observation in the UMAP plots. As shown in the UMAP plots, only scMODAL successfully maintained natural killer (NK) cells, CD4 T cells and CD8 T cells as clearly separated clusters, while in the results of other methods, NK cells were often mixed with effector memory CD8 T (CD8 TEM) cells due to their similarity in RNA modality (Supplementary Fig. 2). Among all compared methods, MaxFuse ranked second in preserving cell population structures but failed to preserve the difference between NK cells and CD8 TEM cells in protein abundance levels (Fig. 2**h**). The other methods also produced less satisfactory integration results. For example, bindSC did not preserve the distinction between NK cells and CD8 TEM cells, and GLUE inaccurately matched these NK cells, CD4 T cells, and CD8 T cells. Portal and Seurat did not integrated CD4 T cells well across modalities (Supplementary Fig. 1).

We also evaluated all methods using a reduced protein panel consisting of the 30 most informative proteins, a typical scenario in single-cell proteomic datasets. Even with this reduced feature set, scMODAL consistently demonstrated superior performance compared to other methods, highlighting its effectiveness in leveraging a limited number of linked features for precise cross-modality integration (Fig. 2**g**).

Using this dataset, we further assessed scMODAL’s capability to predict protein abundance levels for individual cells based on gene expressions. We included MaxFuse and bindSC in this comparison, as they also support cross-modality imputation following integration. Comparing predicted protein abundance levels with ground truth data, scMODAL consistently outperformed these two methods, showing higher correlations (Fig. 2**j**). On average, scMODAL achieved a correlation of 0.53, whereas MaxFuse and bindSC achieved 0.42 and 0.40, representing a relative improvement of 29% and 34% with scMODAL, respectively. In contrast, predictions based solely on protein-coding genes yielded an average correlation of 0.24 with ground truth protein abundances (Fig. 2**k**). Notably, bindSC’s predictions of relative protein abundances were not consistent at the same scale as ground truth measurements, whereas scMODAL reliably recovered protein abundances at the scale close to ground truth with improved correlations, as evidenced by comparisons with bindSC predictions and corresponding protein-coding genes (Fig. 2**i**). These results underscore scMODAL’s proficiency in cross-modality feature imputation even in the absence of paired cells.

Using scMODAL’s predicted features, we were able to generate cells simultaneously measured with features from different modalities, facilitating the identification of feature correlation networks. For instance, we examined the gene-protein correlation network inferred using cells from the RNA modality with imputed proteins. Several genes showed strong correlations with monocyte-enriched proteins, with correlation coefficients exceeding 0.8 or falling below -0.8. These genes are also found to be monocyte-enriched or -depleted genes (Supplementary Fig. 3). As an example, CD64 only exhibited a 0.54 correlation with its coding gene *FCGR1A*, but displayed strong correlations with many other genes (Fig. 2**l**). These findings illustrate scMODAL’s capability to suggest potential gene-protein interactions, thereby elucidating the intricate molecular dynamics within cells through the integration of scRNA-seq and proteomics datasets.

**Figure 3.**
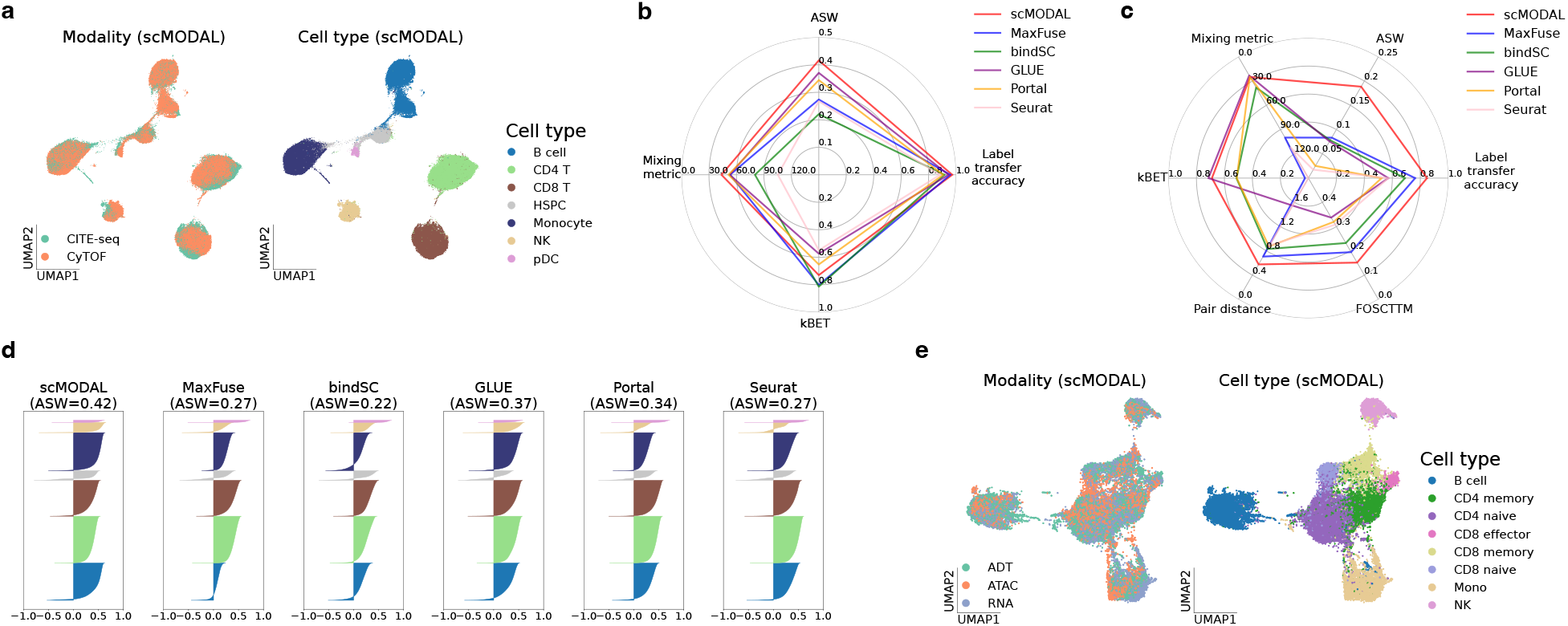
Benchmarking on integration using limited shared features using CITE-seq and CyTOF data and tri-modality integration using TEA-seq data. **a**. UMAP plots of integrated embeddings produced by scMODAL, colored by modalities (left) and cell types (right), for the integration of CITE-seq and CyTOF data. **b**. Quantitative evaluations for the integration of CITE-seq and CyTOF data. **c**. Quantitative evaluations for the tri-modality integration of TEA-seq data. **d**. Bar plots of cell-type silhouette coefficients for individual cells colored by cell types. Larger values indicate better grouping of different cell types. **e**. UMAP plots of integrated embeddings produced by scMODAL, colored by modalities (left) and cell types (right), for the tri-modality integration of TEA-seq data.

### Benchmarking integration of proteomics datasets with limited shared features and tri-modality integration

To further demonstrate scMODAL’s effectiveness, we benchmarked scMODAL against other integration methods in two additional challenging scenarios: one where there are very few shared features, and another where datasets from multiple modalities with varying degrees of shared information are integrated.

In the first scenario, we benchmarked all methods using two human bone marrow single-cell proteomic datasets produced by two different technologies: a sequencing-based CITE-seq dataset [20] and a mass cytometry-based cytometry by time of flight (CyTOF) dataset [35]. In addition to the technical variations between technologies, integrating proteomics datasets from different studies are further complicated by different antibody panels used with only several overlapping markers, providing limited shared information. For instance, for the two datasets we used for benchmarking, the CITE-seq dataset includes 29 protein markers, while the CyTOF dataset includes 32 protein markers, with only 12 markers shared between them.

After applying scMODAL, the datasets were well-mixed in the integrated latent embeddings, as indicated by the UMAP plot and scMODAL’s strong scores in mixing performance (Fig. 3**a, b** and Supplementary Fig. 4). More importantly, the highest label transfer accuracy demonstrated scMODAL’s accuracy in finding correct cell state correspondences across datasets even with limited shared features. Bar plots of cell-type silhouette coefficients revealed that scMODAL produced the best grouping of cell types among all methods, showing its superior performance in preserving biological variations (Fig. 3**d**). We also conducted an ablation study using these datasets to investigate the functionality of each component in scMODAL. This study demonstrated that the adversarial learning objective significantly improves dataset mixing, MNN anchor regularization greatly aids in finding cell state correspondences, and dataset geometric structure regularization helps preserve biological variations by preventing over-correction of cell clusters. More details can be found in the Methods section and Supplementary Fig. 5.

**Figure 4.**
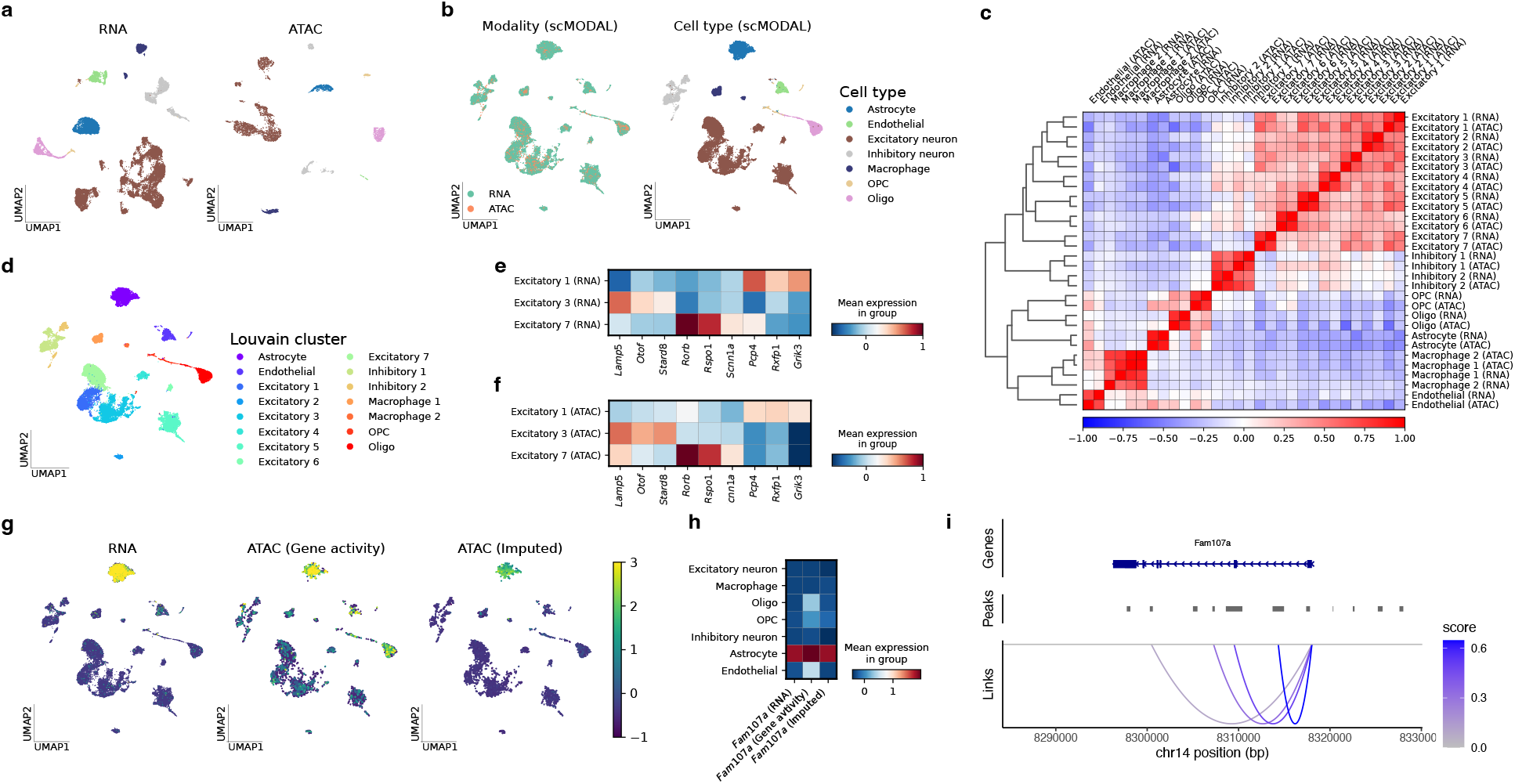
Integration of mouse brain scRNA-seq and scATAC-seq datasets. **a**. UMAP plots of unintegrated scRNA-seq and scATAC-seq datasets, colored by cell types. **b**. UMAP plots of integrated embeddings produced by scMODAL, colored by modalities (left) and cell types (right). **c**. Correlation heatmap of Louvain clusters. **d**. UMAP plots of integrated embeddings produced by scMODAL, colored by Louvain cluster labels. **e, f**. Mouse brain cortical layer marker gene expression levels in the RNA modality (**e**) and gene activity scores in the ATAC modality (**f**) in Louvain clusters 1, 3 and 7. **g, h**. *Fam107a* expression levels in the RNA modality, activity scores in the ATAC modality and imputed expression levels in the ATAC modality shown in UMAP (**g**) and heatmap (**h**). **i**. Gene-peak links inferred using scMODAL’s imputation result.

In the second scenario, we used a human PBMCs dataset profiled by transcription, epitopes, and accessibility sequencing (TEA-seq) [36], which includes 46 protein markers, to evaluate all methods. Specifically, TEA-seq simultaneously measures transcriptomics, epitopes, and chromatin accessibility from cells. This allows us to assess whether an integration method can achieve high-quality tri-modality integration. The challenge in this scenario lies in the higher degree of information sharing between RNA and ATAC modalities compared to RNA and protein [37], requiring integration methods to be flexible and adaptive to handle the heterogeneity in cross-modality gaps.

As shown in the UMAP plots, scMODAL effectively integrated these modalities (Fig. 3**e** and Supplementary Fig. 6), successfully preserving distinct clusters for B cells, T cells, monocytes, and NK cells. scMODAL outperformed or matched the best evaluation metrics among all compared methods, indicating its superior overall integration performance (Fig. 3**c**). Notably, it achieved an RNA-to-ADT label transfer accuracy of 87% and an RNA-to-ATAC label transfer accuracy of 83%, making it the only method to achieve both accuracy scores higher than 70% among all methods (Supplementary Fig. 7). However, not all methods can produce satisfactory results in this tri-modality integration task. Other integration methods had various shortcomings: MaxFuse failed to align different T cell subtypes well, leading to poor scores in mixing performance and cell-state matching accuracy. bindSC, Portal, and Seurat did not adequately maintain separation between cell types, resulting in a loss of biological information and low matching accuracy. Although GLUE showed good alignment of RNA and ATAC modalities, it struggled with ADT modality integration, improperly aligning B cells with monocytes and mismatching different T cell subtypes. The above result highlighted scMODAL’s reliability in handling cross-modality integration tasks with varying degrees of shared variation across datasets.

### Accurate integration of mouse brain scRNA-seq and scATAC-seq datasets enabling peak-gene regularity inference

As a complex organ, the brain contains diverse cell types, including glial cells, endothelial cells, and numerous neuron subtypes. Integrating different single-cell modalities that measure brain cells is crucial for revealing intricate cell type-specific regulatory networks and pathways, as well as studying disease mechanisms such as the Alzheimer’s disease in the brain [10]. However, integrating single-cell brain datasets is challenging because it is difficult to preserve the nuanced difference between neuron subclusters. After validating scMODAL’s effectiveness in cross-modality integration through benchmarking studies, we applied scMODAL to integrate a scRNA-seq dataset [38] and a scATAC-seq obtained from the cortex of mouse brains, demonstrating its ability to achieve accurate integration and facilitate multimodal single-cell analysis in complex organs.

Before integration, different brain cell types in the datasets were annotated according to marker genes (Fig. 4**a**). After scMODAL’s integration, cells of the same cell type were correctly aligned in the latent space (Fig. 4**b**). To validate the integration accuracy in detail, we used the Louvain method [39] to find fine-grained cell type clusters in the integrated cell embedding space (Fig. 4**d**). We identified 15 clusters in total, nine of which corresponded to different neuron subtypes. The clusters were then relabeled according to cell types. For each cluster, there were both RNA modality cells and ATAC modality cells. By comparing the similarity of these clusters using gene expression levels for RNA cells and gene activity scores for ATAC cells, we found that clusters in different modalities assigned the same Louvain cluster label tended to have a higher similarity, as shown by the 2 *×* 2 blocks on the diagonal of the correlation matrix (Fig. 4**c**). This indicates that scMODAL correctly matched corresponding neuron subtypes after integration. We closely examined a major excitatory neuron population formed by excitatory neuron clusters 1, 3, and 7. As shown in Fig. 4**e, f**, many marker genes of mouse brain cortical layers, such as layer 2/3 enriched genes *Lamp5, Otof* and *Stard8*, layer 4 enriched genes *Rorb, Rspo1* and *Scnn1a*, and layer 5/6 enriched genes *Pcp4, Rxfp1* and *Grik3* [40, 41, 42, 43, 44, 45, 46], exhibit consistent differentially expressed patterns in these three clusters in scRNA-seq and scATAC-seq data. Specifically, excitatory neuron clusters 1, 3 and 7 exhibited high layer 4, layer 2/3 and layer 5/6 marker gene expressions, respectively. This result demonstrates that scMODAL correctly aligned detailed cortical neuron cell cluster structures across different modalities. Additionally, by mapping the Louvain cluster labels back to the original space, we found that scMODAL successfully preserved neuron subtype cluster structures contained in the original datasets. For example, excitatory neuron clusters 2, 4, 5 and 6 form isolated clusters in the unintegrated datasets, unconnected from the major excitatory neuron population formed by clusters 1, 3 and 7 (Supplementary Fig. 8). This pattern is well-preserved after scMODAL integration, demonstrating its ability to maintain the subtle similarities and differences among neuronal subtypes.

For comparison, we also applied MaxFuse, which ranked second in integration accuracy in our benchmarking studies, and GLUE, a state-of-the-art method for integrating scRNA-seq and scATAC-seq data, to integrate these two datasets. Comparing to scMODAL, these two methods produced less accurate integration results in terms of finding cell-state matching (Supplementary Fig. 9). For example in MaxFuse’s integration, a cluster of excitatory neurons and a cluster of inhibitory neurons were incorrectly aligned with each other.

Using scMODAL’s integration result, we performed gene expression imputation for the scATAC-seq data to generate virtual cells with simultaneous measurement of gene expression and chromatin accessibility. Interestingly, we found the imputation results for some genes align with the gene expression patterns observed in scRNA-seq data but differ from the gene activity scores in scATAC-seq data. This discrepancy arises because gene score prediction methods using scATAC-seq data often assume that chromatin accessibility within the gene locus or nearby regions consistently contributes to improving the gene expression level, which may not reflect the true regulatory mechanisms. For example, consider the astrocyte marker gene *Fam107a* [47]. In the scRNA-seq dataset, *Fam107a* shows high expression exclusively in astrocytes, but it is depleted in other cell types (Fig. 4**g, h**). However, the gene activity scores produced by Signac [12] infer *Fam107a* expression in oligodendrocytes, oligodendrocyte progenitor cells (OPC) and endothelial cells, likely due to chromatin accessibility peaks detected near the *Fam107a* gene (Supplementary Fig. 10). In contrast, scMODAL’s gene imputation results show *Fam107a* expression patterns that more closely resemble the scRNA-seq data, with clear enrichment only in astrocytes. We further investigated potential *cis*-regulatory interactions by calculating the correlation coefficients between the imputed gene expression and the accessibility of each peak within a 10kb distance from *Fam107a* [48]. As shown in Fig. 4**i**, based on scMODAL’s imputation, only peaks highly accessible in astrocytes were inferred to be associated with *Fam107a*, providing a reduced candidate peak set that could potentially regulate the gene expression level of *Fam107a* in the brain. The above analysis demonstrated scMODAL’s ability to provide insights to regulatory signatures using unpaired multi-omics single-cell datasets.

### Integration of CODEX and scRNA-seq facilitating spatial structure identification of B cell follicles in tonsil

As a recently developed technology, co-detection by indexing (CODEX) enables highly multi-plexed and spatially resolved profiling of proteins within tissue sections at single-cell resolution [6]. This technique has been widely applied to study diverse immune microenvironments, such as those in lymph nodes [49] and tumors [50]. However, to accurately characterize a specific immune microenvironment using single-cell spatial proteomics, it is essential to have a well-designed protein panel targeting specific cell types of interest. In this section, we show how integrating single-cell spatial proteomics data with scRNA-seq data using scMODAL can improve the spatial characterization of the microenvironment, even when the protein panel is not fully comprehensive. This integration is illustrated using a human tonsil CODEX dataset including 44 protein markers [51] and a corresponding tonsil scRNA-seq dataset [52].

Using the CODEX tonsil section with the original cell-type annotation, we identified B cell follicle structures organized around interfollicular regions rich in T cells (Fig. 5**a**). In these interfollicular regions, T cells interact with B cells, facilitating the formation of germinal centers within the B cell follicles [53]. Within these germinal centers, mature B cells undergo activation, proliferation, differentiation, and diversify their antibody genes through somatic hypermutation [54, 55]. However, the CODEX data protein panel did not include proliferation-associated markers such as Ki67, which is crucial for identifying proliferating germinal center B cells [56], or marginal zone B cell markers like CD22 and CD40 [57]. This limitation makes it challenging to identify B cell subtypes and fully characterize the structure of B cell follicles in the tonsil section (Supplementary Fig. 11).

**Figure 5.**
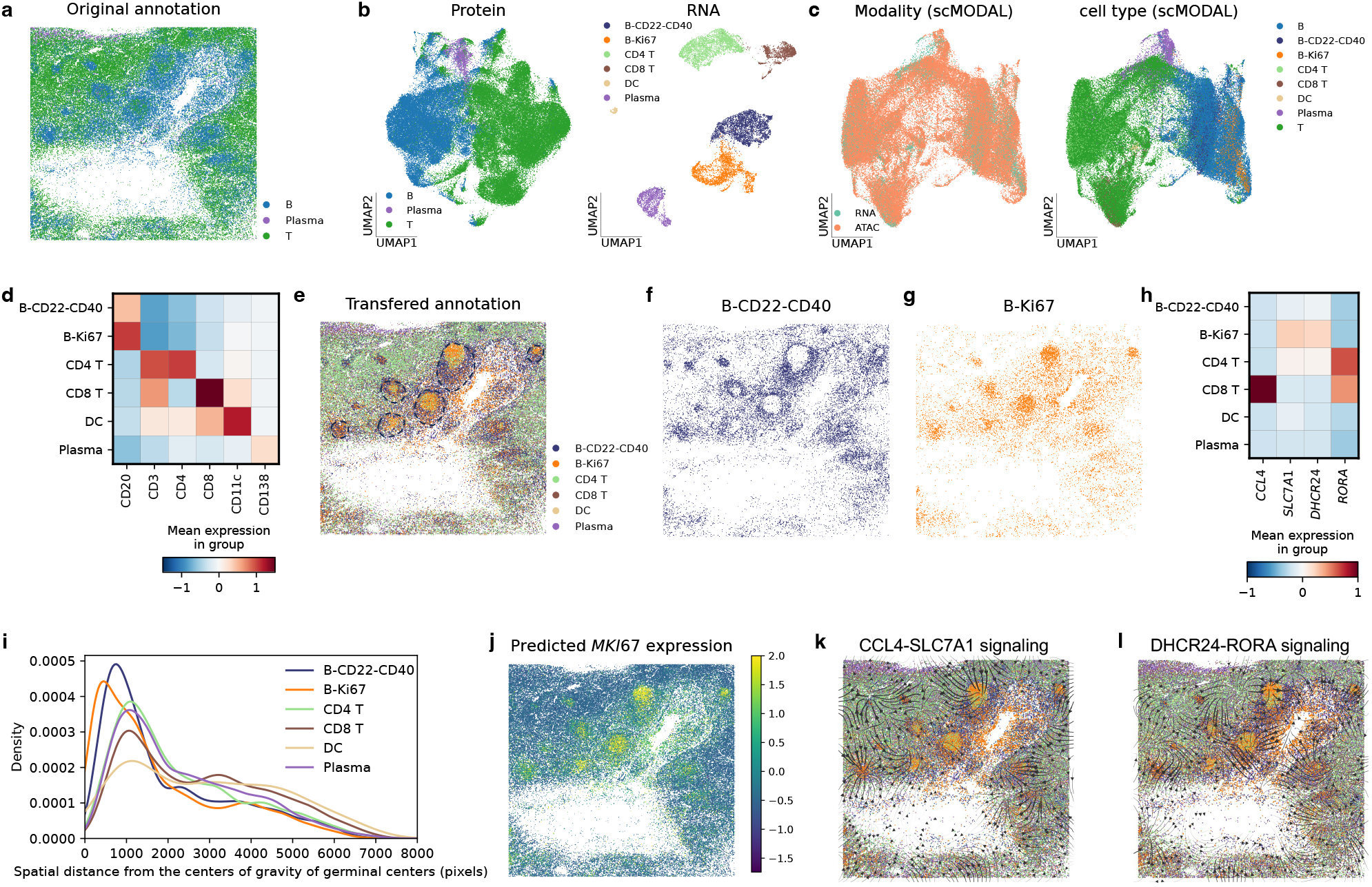
Integration of human tonsil CODEX and scRNA-seq datasets. **a**. The CODEX human tonsil section with the original cell-type annotation. **b**. UMAP plots of unintegrated CODEX and scRNA-seq datasets, colored by cell types. **c**. UMAP plots of integrated embeddings produced by scMODAL, colored by modalities (left) and cell types (right). **d**. Cell-type markers in the CODEX dataset in cell types identified with transferred labels. **e**. The CODEX human tonsil section with transferred cell-type annotation. Dashed circles indicate regions showing clear B cell follicle structures. **f, g**. Spatial distributions of B-CD22-CD40 (**f**) and B-Ki67 cells (**g**). **h**. Predicted *DHCR24, RORA, CCL4* and *SLC7A1* expression levels in the CODEX dataset in cell types identified with transferred labels. **i**. Distributions of spatial distances between cells of different cell types and the centers of gravity of germinal centers. **j**. Predicted *MKI67* spatial expression pattern in the CODEX dataset. **k, l**. The spatial signaling directions of CCL4-SLC7A1 signaling (**k**) and DHCR24-RORA signaling (**l**) pathways inferred by COMMOT.

Unlike the CODEX dataset, the scRNA-seq dataset clearly distinguishes between germinal center B cells (B-Ki67) and marginal zone B cells (B-CD22-CD40), as well as between CD4 and CD8 T cells (Fig. 5**b**). By integrating the CODEX and scRNA-seq tonsil datasets using scMODAL (Fig. 5**c**), we successfully transferred cell-type labels from the scRNA-seq data to the CODEX data. After this label transfer, we validated the results using the available protein panel (Fig. 5**d**). The protein abundance confirmed that CD4 and CD8 T cells identified by scMODAL in the CODEX data were specifically enriched for CD4 and CD8, respectively, while all expressed the T cell marker CD3. Additionally, following the label transfer, B cells, dendritic cells (DCs), and plasma cells were enriched for their corresponding markers—CD20, CD11c, and CD138, respectively. Importantly, the cell types identified with the transferred annotation displayed distinct spatial distribution patterns (Fig. 5**e**). For the B-CD22-CD40 subtype, we observed that these cells formed several hollow circles in the outer regions of B cell follicles, indicating the presence of marginal zones (Fig. 5**f**). Additionally, B-Ki67 cells were concentrated within the circles formed by marginal zone B cells, marking the spatial locations of germinal centers (Fig. 5**g**). Together, these two B cell subtypes, identified through transferred cell-type labels, revealed the spatial organization of B cell follicle structures. Using DBSCAN [58], we identified six germinal centers using the spatial distribution of B-Ki67 cells (Supplementary Fig. 12) and calculated the minimum distance between each cell and the center of gravity of any germinal center. The distance distributions confirmed expected spatial patterns, with B-Ki67, B-CD22-CD40, and other cells showing increasing distances from the germinal centers (Fig. 5**i**).

The gene expression levels predicted by scMODAL further enhance the spatial characterization of the tonsil section. For example, we imputed the expression level of *MKI67*, the gene encoding Ki67 (Fig. 5**j**). Although Ki67 abundance was not measured in the original CODEX dataset, the imputed *MKI67* expression accurately captured B cell dynamics. Specifically, imputed *MKI67* showed high expression in germinal centers with a decreasing gradient from the inner to outer B cell follicles, reflecting the spatial specificity of B cell proliferation. Leveraging scMODAL’s imputed gene expression levels in the CODEX tonsil section, we further applied COMMOT [59] to analyze cell-cell communication within the tonsil immune microenvironment, using ligand-receptor information from the CellPhoneDB database [60]. For instance, the CCL4-SLC7A1 interaction, which has been used to study immune cell communication pathways [61], was explored. In the tonsil scRNA-seq dataset, *CCL4* was enriched in CD8 T cells, while *SLC7A1* was enriched in B-Ki67 cells (Fig. 5**h**). This interaction was identified between germinal center B cells and interfollicular T cells, suggesting a potential B cell-T cell communication pathway in the immune response (Fig. 5**k**). Additionally, we identified other spatial cell-cell communication pathways, such as DHCR24-RORA signaling between B cells and T cells (Fig. 5**l**). These findings demonstrate scMODAL’s capability in facilitating spatial multi-omics analysis.

## Discussion

In this study, we introduced scMODAL, a novel deep learning framework designed for the integration of single-cell multi-omics data, specifically addressing the challenges associated with datasets that have limited numbers of known correlated features. Our results demonstrate that scMODAL effectively aligns cell embeddings across different modalities, preserves the biological variation, and accurately identifies cell subpopulations. Moreover, scMODAL excels in tasks such as cross-modality feature imputation and inferring feature relationships, which are critical for understanding the underlying cellular processes.

Compared to existing methods such as MaxFuse, bindSC, GLUE, Portal, and Seurat, scMODAL offers distinct advantages. While methods like MaxFuse and bindSC have shown efficacy in integrating modalities with weak relationships, they rely on linear projections that may not fully capture the complex, nonlinear nature of unwanted variations present in multi-omics datasets. Additionally, methods designed for integrating scRNA-seq and scATAC-seq datasets, such as GLUE, might overestimate the shared information across modalities that are only weakly connected, such as transcriptomics and proteomics, leading to inaccurate integration results in this setting. scMODAL addresses these limitations by employing nonlinear neural networks and GANs to align cell embeddings, ensuring that the integration process retains the intrinsic biological structure of the data. Notably, scMODAL’s use of MNN pairs as anchors during the integration process plays a crucial role in improving the accuracy of the integration, even when the number of linked features is limited. The benchmarking results on a collection of datasets in different scenarios highlight scMODAL’s superiority in mixing cell distributions, maintaining cell-type separations, and accurately matching corresponding cell states across modalities.

The ability of scMODAL to preserve biological variation while integrating multi-omics data has significant implications for the study of complex cellular processes. For instance, its capacity to accurately identify cell subpopulations that were not distinguishable with individual modalities suggests that scMODAL could be instrumental in uncovering new cell types or states. Additionally, the feature imputation capabilities of scMODAL could facilitate the discovery of novel gene regulatory networks and pathways that are otherwise obscured in single-modality analyses. The gene-protein and gene-peak link inference, along with the discovery of spatial cell-cell communication patterns using scMODAL’s imputation results, exemplify the practical utility of this functionality.

Despite its advantages, scMODAL has certain limitations. For example, the reliance on known linked features for integration, although effective, may limit its applicability to scenarios where such features are not well-characterized or absent. Future work could explore the incorporation of unsupervised learning techniques to identify potential links between modalities, thereby broadening the applicability of scMODAL.

In conclusion, scMODAL represents a significant advancement in the field of single-cell multi-omics data integration. By leveraging deep learning techniques, it addresses the critical challenges of cross-modality integration, offering a robust tool for researchers to explore the complex interplay between different cellular components. As single-cell technologies continue to evolve, frameworks like scMODAL will be indispensable in translating multi-omics data into actionable biological insights.

## Methods

### The model of scMODAL

Let 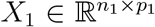 and 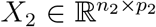 be the matrices representing single-cell features from two different modalities. As features in the two modalities are usually not shared, prior knowledge about the cross-modality feature relationships is required for finding correspondence between modalities. We construct a new pair of matrices 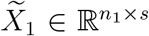 and 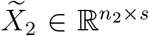, with *s* pairs of features likely to positively correlate with each other based on prior information. For the integration of proteomic and scRNA-seq data, we let each pair be the abundance of a protein and the expression level of its coding gene. When integrating scRNA-seq and scATAC-seq data, we used gene expression levels and gene activity scores of shared highly variable genes as feature pairs.

#### Aligning different modalities using generative adversarial learning

To integrate datasets while preserving biological information contained in all highly variable features, we introduce a shared latent space *Z* and encode information into *Z* with neural networks. Denote the encoders as *E*_1_(*·*) and *E*_2_(*·*), our goal is to integrate the cell embedding distributions *E*_1_(**x**_1_) and *E*_2_(**x**_2_) in *Z*, where **x**_1_ and **x**_2_ represent cells from *X*_1_ and *X*_2_, respectively. We apply generative adversarial learning to align the empirical distributions of *E*_1_(**x**_1_) and *E*_2_(**x**_2_) in *Z*, borrowing the idea from Generative Adversarial Networks (GANs) [30]. Specifically, we use an auxiliary network *D*(*·*) : *Z →* (0, 1) as the discriminator to distinguish cell embeddings from two datasets by maximizing the following objective:

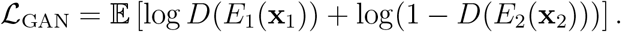

The encoders are trained against the discriminator by minimizing *ℒ*_GAN_, which is equivalent to minimizing the Jensen-Shannon (JS) divergence between the distributions of *E*_1_(**x**_1_) and *E*_2_(**x**_2_) [30]. This process is represented by the minimax optimization formula min_*E*_1,*E*2 max_*D*_ *ℒ*_GAN_.

#### Regularization for within- and cross-domain autoencoding consistency

Two decoders, denoted as *G*_1_(*·*) and *G*_2_(*·*), are introduced and trained together to ensure within- and cross-domain autoencoding consistency by minimizing the autoencoder loss:

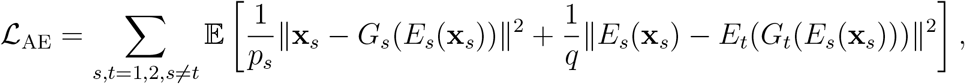

where *q* is the dimensionality of *Z*.

#### Regularization for aligning anchors with prior feature linkage information

Using generative adversarial learning to align distributions without constraints can result in incorrect matching of cell populations. To learn accurate integration results, we utilize similarity information in linked features to guide integration. Specifically, during minibatch training with two minibatches from two modalities, for each cell in one minibatch, we find the *k*-nearest neighborhoods in cells in another minibatch by comparing the angle distance between corresponding linked features in 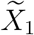 and 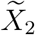, and vice versa. This procedure gives us mutual nearest neighborhood pairs 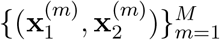, serving as anchors for integration. For these pairs, we let their embeddings to be close to each other by minimizing the objective:

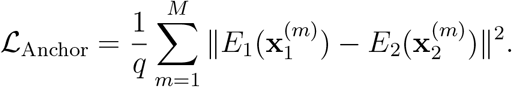

#### Regularization for data structure preservation

To avoid loss of information contained in dataset-unique features, we propose to preserve the geometric structure of each dataset by regularizing the geometric representations of cells. To be specific, for a cell 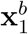 from *X*_1_ in a minibatch, we calculate the Gaussian kernel distance with all cells in the batch as its geometric representation:

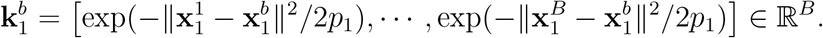

The geometric representation is also calculated for the cell representation of 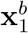 in *Z* as

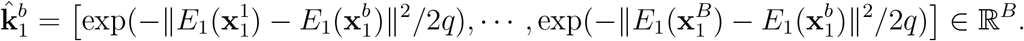

Similarly we define the geometric representations of cells from *X*_2_ in the other minibatch. The geometric representation of a cell indicates its relative distance from other cells computed with all variable features. We use a geometric structure regularization to preserve this information by minimizing the objective.

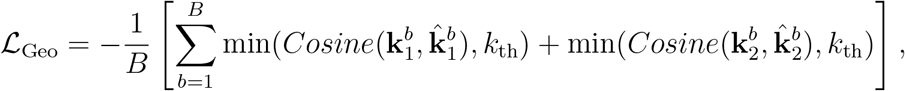

where *k*_th_ = 0.975 is a fixed threshold.

#### Training procedure

To integrate cross-modality datasets with correctly matched cell states while preserving important biological variation, we train the networks by considering the generative adversarial learning objective and other regularizers jointly in the following mini-max optimization formula

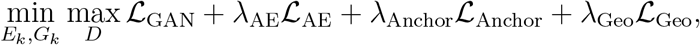

where *λ*_AE_, *λ*_Anchor_ and *λ*_Geo_ are coefficients for the regularizers. During training, the neural networks in scMODAL are updated iteratively to solve the mini-max problem. Once the training is finished, cell embeddings in *Z* serve as integrated representations for further downstream tasks. Besides, *G*_2_(*E*_1_(*·*)) and *G*_1_(*E*_2_(*·*)) can be used to predict unmeasured features across modalities.

### Analysis details

#### Integration of multiple datasets

Benefiting from scalable neural network training, scMODAL can also be used for integrating multiple multi-omics datasets. When there are more than two datasets to be integrated (denoted as 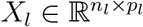, *l* = 1, 2, *· · ·, L*), scMODAL handels the integration task by introducing *L −* 1 discriminators to align dataset pairs (*X*_*l*_, *X*_*l*+1_), *l* = 1, 2, *· · ·, L −* 1 in the latent space *Z*. The regularizers for cross-domain autoencoding consistency and aligning anchors with prior feature linkage information are also extended accordingly for dataset pairs (*X*_*l*_, *X*_*l*+1_).

For integrating multiple modalities, we found that scMODAL is robust to the order in which modalities are processed. For instance, in the tri-modality integration task using the TEA-seq dataset, we explored the relationships between ADT and RNA, as well as between RNA and ATAC, setting the integration order as (*X*_1_, *X*_2_, *X*_3_) = (*X*_ADT_, *X*_RNA_, *X*_ATAC_). We also evaluated scMODAL’s performance with orders (*X*_1_, *X*_2_, *X*_3_) = (*X*_ADT_, *X*_ATAC_, *X*_RNA_) and (*X*_1_, *X*_2_, *X*_3_) = (*X*_RNA_, *X*_ADT_, *X*_ATAC_), with different modalities serving as the bridge between the other two. Overall, scMODAL demonstrated consistently strong integration performance regardless of the modality order (Supplementary Fig. 13).

#### Model training details

scMODAL employs the Adam optimizer [62] for stochastic optimization during model training. By default, the batch size is set to *B* = 500. The optimization process runs for 10,000 iterations with a learning rate of *lr* = 0.001, coefficients for running averages (*β*_1_, *β*_2_) = (0.9, 0.999), and a weight decay parameter of *λ* = 0.001 across all networks. The latent space dimensionality is set to *q* = 20 and the neighborhood size is set to *k* = 30 for identifying MNNs. The regularization parameters are *λ*_AE_ = 10.0, *λ*_Anchor_ = 1.0, and *λ*_Geo_ = 1.0.

#### Computational time and memory usage

We evaluated the computational time and memory usage of all methods using the CITE-seq PBMC dataset [31] with different sample sizes. For the benchmarking of computational time and memory usage, we applied all methods on the same Linux server with Intel Xeon Gold 5222 CPUs. For methods that require GPUs including scMODAL, GLUE and Portal, a single NVIDIA RTX 5000 GPU was used in all the experiments. To only focus on the integration algorithms, we only recorded the running time and memory usage after standard data preprocessing such as normalization, scaling and dimension reduction. As illustrated in Supplementary Fig. 14, bindSC and Seurat were unable to complete the integration of datasets with 200,000 and 300,000 cells, respectively, due to their peak memory usage exceeding the 160 GB limit. Unlike these two methods, scMODAL demonstrates efficient memory usage, allowing it to handle large datasets without exceeding memory limits. Moreover, as dataset size increases, scMODAL demonstrates faster running times compared to MaxFuse and GLUE, highlighting its training efficiency.

#### Ablation study

We investigated the functionalities of different components in scMODAL’s model using the CITE-seq and the CyTOF human bone marrow datasets. As shown in Supplementary Fig. 5, we observed that removing the GAN objective from the loss function led to less well-mixed cell distributions, as evidenced by a higher mixing metric and a lower kBET metric. Additionally, removing the regularization for autoencoding consistency resulted in less accurate cell-state matching, reflected in a decrease in label transfer accuracy. This also led to poorer preservation of biological variation, as indicated by a lower ASW score. When the regularization for aligning MNN anchors was removed, nearly all cell states were incorrectly matched, with label transfer accuracy approaching 0, indicating that cells were aligned with others of different cell type labels. Furthermore, removing the regularization for data structure preservation caused a decrease in the ASW score, suggesting a decline in the preservation of cell-type cluster information.

#### Evaluation metrics

We used the mixing metric [20] and *k*-nearest-neighbor batch-effect test (kBET) [33] to assess the ability of unwanted variation removal. Besides, we used the average silhouette width (ASW) to evaluate the preservation of biological variation, and label transfer accuracy, pair distance, and fraction of samples closer than true match (FOSCTTM) [34] to measure the correctness of cell-state matching in the integration results.

#### Mixing metric

For each cell, the rank in its *k* = 300 neighborhood corresponding to the fifth neighbor in each dataset is calculated. The mixing metric is then obtained by taking the median of the ranks over all datasets and then taking the average over all cells. A lower mixing metric indicates better mixing of the datasets.

#### kBET

kBET uses a Pearson’s *χ*^2^-based test to evaluate whether the distribution of batch labels in the neighborhood of a cell matches the overall distribution of batch labels. In our experiments, we ran 100 replicates, each with 1,000 randomly selected samples, and we used the median of the average acceptance rates as the final output. A higher kBET indicates better mixing of the datasets.

#### ASW

For each cell, its silhouette width is defined as (*b − a*)*/* max(*a, b*), where *a* represents the mean distance between the cell and other cells within the same cluster, and *b* represents the mean distance between the cell and other cells from the nearest cluster that the cell does not belong to. The ASW score is then the average of silhouette widths over all cells. A higher ASW indicates a better preservation of clustering structures in the integration results.

#### Label transfer accuracy

In the integrated cell embedding space, we transfer labels from one dataset to another dataset based on the nearest neighbor using the euclidean distance. Then we evaluate the ratio of correct transferred labels as the label transfer accuracy. A higher label transfer accuracy indicates a more accurate matching of corresponding cell states.

#### Pair distance

This metric is evaluated with single-cell multi-omics datasets with simultaneously measured features from different modalities. Given the ground truth pairs of embeddings from different modalities, say 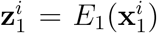and 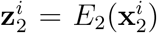, the pair distance is a relative distance defined as 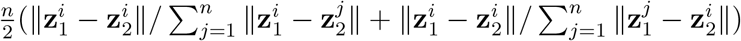, where *n* is the total number of cells. The final score is then the average of pair distances over all cells. A lower pair distance indicates the ground truth pairs from different modalities are better matched after integration.

#### FOSCTTM

Given the ground truth pairs of embeddings 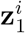 and 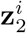, FOSCTTM is computed as 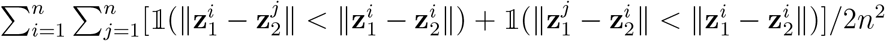. A lower FOSCTTM means that the ground truth pairs have closer distances, indicating a better integration result.

## Data availability

All data used in this work are publicly available through online sources.

- Human PBMC CITE-seq dataset [31] (GSE164378).
- Human bone marrow CITE-seq dataset [20] (GSE128639).
- Human bone marrow CyTOF dataset [35] (https://github.com/lmweber/benchmark-data-Levine-32-dim).
- Human PBMC TEA-seq dataset [36] (GSE158013).
- Mouse brain scRNA-seq dataset profiled by 10x Genomics [38] (http://mousebrain.org/adolescent/downloads.html).
- Mouse brain scATAC-seq dataset profiled 10x Genomics (http://cf.10xgenomics.com/samples/cell-atac/1.1.0/atac_v1_adult_brain_fresh_5k/atac_v1_adult_brain_fresh_5k_filtered_peak_bc_matrix.h5).
- Human tonsil CODEX dataset [51] (https://datadryad.org/stash/share/1OQtxew0Unh3iAdP-ELew-ctwuPTBz6Oy8uuyxqliZk).
- Human tonsil scRNA-seq dataset [52] (GSE165860).

## Code availability

scMODAL software is available at https://github.com/gefeiwang/scMODAL.

## Acknowledgements

This work was supported in part by NIH grants R01 GM134005, U24 HG012108, P50 CA196530, R01 AG068191, RF1 AG081413 and R01 EB034720.

## References

[1] Tang, F. et al. mRNA-Seq whole-transcriptome analysis of a single cell. Nature Methods 6, 377–382 (2009).

[2] Buenrostro, J. D. et al. Single-cell chromatin accessibility reveals principles of regulatory variation. Nature 523, 486–490 (2015).

[3] Cusanovich, D. A. et al. Multiplex single-cell profiling of chromatin accessibility by combinatorial cellular indexing. Science 348, 910–914 (2015).

[4] Bendall, S. C. et al. Single-cell mass cytometry of differential immune and drug responses across a human hematopoietic continuum. Science 332, 687–696 (2011).

[5] Stoeckius, M. et al. Simultaneous epitope and transcriptome measurement in single cells. Nature Methods 14, 865–868 (2017).

[6] Goltsev, Y. et al. Deep profiling of mouse splenic architecture with CODEX multiplexed imaging. Cell 174, 968–981 (2018).

[7] Villani, A.-C. et al. Single-cell RNA-seq reveals new types of human blood dendritic cells, monocytes, and progenitors. Science 356, eaah4573 (2017).

[8] Hickey, J. W. et al. Organization of the human intestine at single-cell resolution. Nature 619, 572–584 (2023).

[9] Frangieh, C. J. et al. Multimodal pooled Perturb-CITE-seq screens in patient models define mechanisms of cancer immune evasion. Nature Genetics 53, 332–341 (2021).

[10] Xiong, X. et al. Epigenomic dissection of Alzheimer’s disease pinpoints causal variants and reveals epigenome erosion. Cell 186, 4422–4437.e21 (2023).

[11] Argelaguet, R., Cuomo, A. S., Stegle, O. & Marioni, J. C. Computational principles and challenges in single-cell data integration. Nature Biotechnology 39, 1202–1215 (2021).

[12] Stuart, T., Srivastava, A., Madad, S., Lareau, C. A. & Satija, R. Single-cell chromatin state analysis with Signac. Nature Methods 18, 1333–1341 (2021).

[13] Granja, J. M. et al. ArchR is a scalable software package for integrative single-cell chromatin accessibility analysis. Nature Genetics 53, 403–411 (2021).

[14] Schwanhäusser, B. et al. Global quantification of mammalian gene expression control. Nature 473, 337–342 (2011).

[15] Vogel, C. & Marcotte, E. M. Insights into the regulation of protein abundance from proteomic and transcriptomic analyses. Nature Reviews Genetics 13, 227–232 (2012).

[16] Liu, Y., Beyer, A. & Aebersold, R. On the dependency of cellular protein levels on mRNA abundance. Cell 165, 535–550 (2016).

[17] Battle, A. et al. Impact of regulatory variation from RNA to protein. Science 347, 664–667 (2015).

[18] Spitzer, M. H. & Nolan, G. P. Mass cytometry: single cells, many features. Cell 165, 780–791 (2016).

[19] Wang, Y. et al. Spatial transcriptomics: Technologies, applications and experimental considerations. Genomics 110671 (2023).

[20] Stuart, T. et al. Comprehensive integration of single-cell data. Cell 177, 1888–1902.e21 (2019).

[21] Korsunsky, I. et al. Fast, sensitive and accurate integration of single-cell data with Harmony. Nature Methods 16, 1289–1296 (2019).

[22] Lopez, R., Regier, J., Cole, M. B., Jordan, M. I. & Yosef, N. Deep generative modeling for single-cell transcriptomics. Nature Methods 15, 1053–1058 (2018).

[23] Zhao, J. et al. Adversarial domain translation networks for integrating large-scale atlas-level single-cell datasets. Nature Computational Science 2, 317–330 (2022).

[24] Cao, Z.-J. & Gao, G. Multi-omics single-cell data integration and regulatory inference with graph-linked embedding. Nature Biotechnology 40, 1458–1466 (2022).

[25] Dou, J. et al. Bi-order multimodal integration of single-cell data. Genome Biology 23, 112 (2022).

[26] Chen, S. et al. Integration of spatial and single-cell data across modalities with weakly linked features. Nature Biotechnology 42, 1096–1106 (2024).

[27] Haghverdi, L., Lun, A. T., Morgan, M. D. & Marioni, J. C. Batch effects in single-cell RNA-sequencing data are corrected by matching mutual nearest neighbors. Nature Biotechnology 36, 421–427 (2018).

[28] Luecken, M. D. et al. Benchmarking atlas-level data integration in single-cell genomics. Nature Methods 19, 41–50 (2022).

[29] Tran, H. T. N. et al. A benchmark of batch-effect correction methods for single-cell RNA sequencing data. Genome Biology 21, 12 (2020).

[30] Goodfellow, I. et al. Generative adversarial nets. In Advances in Neural Information Processing Systems, 2672–2680 (2014).

[31] Hao, Y. et al. Integrated analysis of multimodal single-cell data. Cell 184, 3573–3587.e29 (2021).

[32] McInnes, L., Healy, J., Saul, N. & Grossberger, L. UMAP: Uniform manifold approximation and projection. The Journal of Open Source Software 3, 861 (2018).

[33] Büttner, M., Miao, Z., Wolf, F. A., Teichmann, S. A. & Theis, F. J. A test metric for assessing single-cell RNA-seq batch correction. Nature Methods 16, 43–49 (2019).

[34] Singh, R. et al. Unsupervised manifold alignment for single-cell multi-omics data. In Proceedings of the 11th ACM International Conference on Bioinformatics, Computational Biology and Health Informatics, BCB ‘20 (2020).

[35] Levine, J. H. et al. Data-driven phenotypic dissection of AML reveals progenitor-like cells that correlate with prognosis. Cell 162, 184–197 (2015).

[36] Swanson, E. et al. Simultaneous trimodal single-cell measurement of transcripts, epitopes, and chromatin accessibility using TEA-seq. Elife 10, e63632 (2021).

[37] Lin, K. Z. & Zhang, N. R. Quantifying common and distinct information in single-cell multimodal data with Tilted Canonical Correlation Analysis. Proceedings of the National Academy of Sciences 120, e2303647120 (2023).

[38] Zeisel, A. et al. Molecular architecture of the mouse nervous system. Cell 174, 999–1014.e22 (2018).

[39] Blondel, V. D., Guillaume, J.-L., Lambiotte, R. & Lefebvre, E. Fast unfolding of communities in large networks. Journal of Statistical Mechanics: Theory and Experiment 2008, P10008 (2008).

[40] Tiveron, M.-C. et al. LAMP5 fine-tunes GABAergic synaptic transmission in defined circuits of the mouse brain. PLOS ONE 11, e0157052 (2016).

[41] Tasic, B. et al. Adult mouse cortical cell taxonomy revealed by single cell transcriptomics. Nature Neuroscience 19, 335–346 (2016).

[42] Shrestha, P., Mousa, A. & Heintz, N. Layer 2/3 pyramidal cells in the medial prefrontal cortex moderate stress induced depressive behaviors. Elife 4, e08752 (2015).

[43] Weed, N. et al. Identification of genetic markers for cortical areas using a random forest classification routine and the Allen Mouse Brain Atlas. PLOS ONE 14, e0212898 (2019).

[44] Bulfone, A. et al. Pcp4l1, a novel gene encoding a Pcp4-like polypeptide, is expressed in specific domains of the developing brain. Gene Expression Patterns 4, 297–301 (2004).

[45] Belgard, T. G. et al. A transcriptomic atlas of mouse neocortical layers. Neuron 71, 605–616 (2011).

[46] Selvakumar, P. et al. Structural and compositional diversity in the kainate receptor family. Cell Reports 37, 109891 (2021).

[47] Batiuk, M. Y. et al. Identification of region-specific astrocyte subtypes at single cell resolution. Nature Communications 11, 1220 (2020).

[48] Ma, S. et al. Chromatin potential identified by shared single-cell profiling of RNA and chromatin. Cell 183, 1103–1116.e20 (2020).

[49] Roider, T. et al. Multimodal and spatially resolved profiling identifies distinct patterns of T cell infiltration in nodal B cell lymphoma entities. Nature Cell Biology 26, 478–489 (2024).

[50] Quek, C. et al. Single-cell spatial multiomics reveals tumor microenvironment vulnerabilities in cancer resistance to immunotherapy. Cell Reports 43, 114392 (2024).

[51] Brbić, M. et al. Annotation of spatially resolved single-cell data with STELLAR. Nature Methods 19, 1411–1418 (2022).

[52] King, H. W. et al. Integrated single-cell transcriptomics and epigenomics reveals strong germinal center–associated etiology of autoimmune risk loci. Science Immunology 6, eabh3768 (2021).

[53] Kennedy, D. E. & Clark, M. R. Compartments and connections within the germinal center. Frontiers in Immunology 12, 659151 (2021).

[54] Klein, U. & Dalla-Favera, R. Germinal centres: role in B-cell physiology and malignancy. Nature Reviews Immunology 8, 22–33 (2008).

[55] De Silva, N. S. & Klein, U. Dynamics of B cells in germinal centres. Nature Reviews Immunology 15, 137–148 (2015).

[56] Roughan, J. E., Torgbor, C. & Thorley-Lawson, D. A. Germinal center B cells latently infected with Epstein-Barr virus proliferate extensively but do not increase in number. Journal of Virology 84, 1158–1168 (2010).

[57] Demberg, T. et al. Loss of marginal zone B-cells in SHIVSF162P4 challenged rhesus macaques despite control of viremia to low or undetectable levels in chronic infection. Virology 484, 323–333 (2015).

[58] Ester, M., Kriegel, H.-P., Sander, J., Xu, X. et al. A density-based algorithm for discovering clusters in large spatial databases with noise. In kdd, vol. 96, 226–231 (1996).

[59] Cang, Z. et al. Screening cell–cell communication in spatial transcriptomics via collective optimal transport. Nature Methods 20, 218–228 (2023).

[60] Garcia-Alonso, L. et al. Single-cell roadmap of human gonadal development. Nature 607, 540–547 (2022).

[61] Lu, C. et al. Single-cell transcriptome analysis and protein profiling reveal broad immune system activation in IgG4-related disease. JCI Insight 8 (2023).

[62] Kingma, D. P. & Ba, J. Adam: A method for stochastic optimization. In International Conference on Learning Representations (2015).

