## Supplementary Information for "scMODAL: A general deep learning framework for comprehensive single-cell multi-omics data alignment with feature links"

### Supplementary Figures

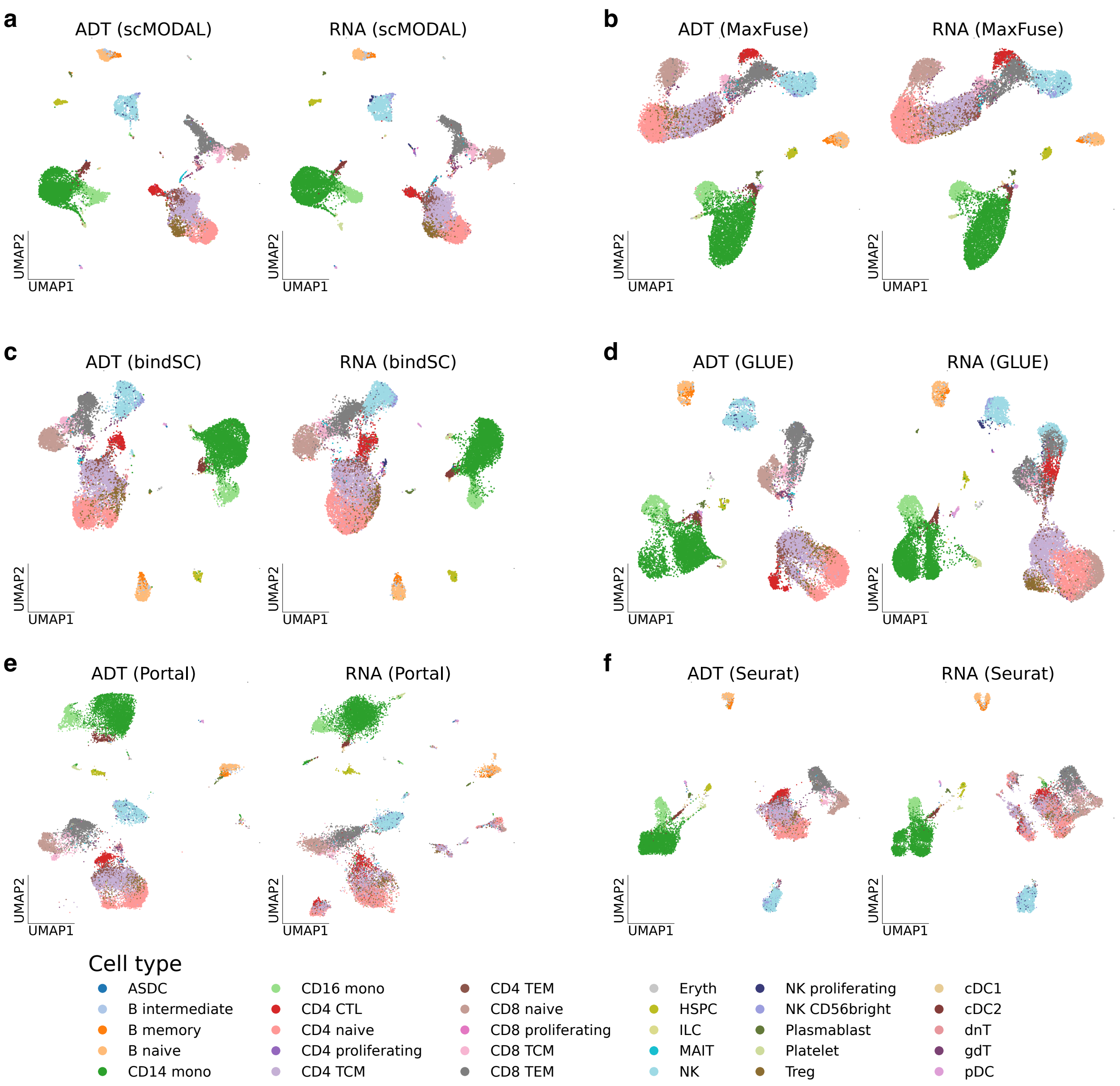


Supplementary Fig. 1: **Comparison of integration methods based on the CITE-seq human PBMC data [1]. a-f.** UMAP [2] plots of integrated embeddings produced by scMODAL (**a**), MaxFuse [3] (**b**), bindSC [4] (**c**), GLUE [5] (**d**), Portal [6] (**e**) and Seurat [7] (**f**), colored by cell types.

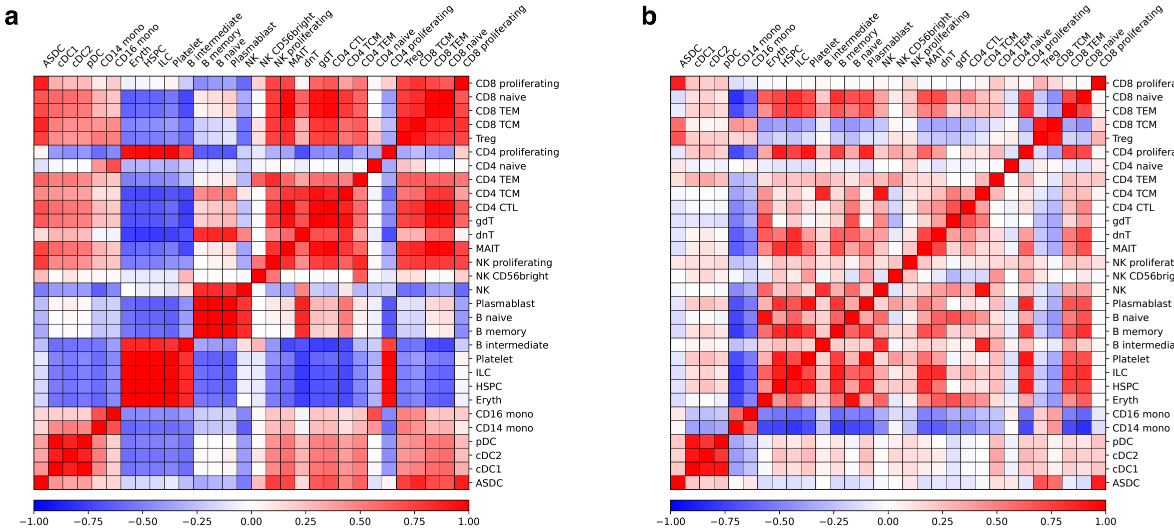


Supplementary Fig. 2: **Correlation heatmaps of different cell types in the CITE-seq human PBMC data. a**, **b**. Correlation heatmaps calculated using RNA features (**a**) and ADT features (**b**).

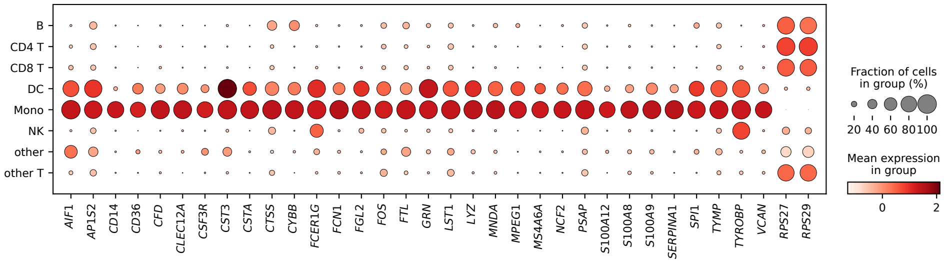


Supplementary Fig. 3: **Expression levels of genes positively or negatively correlated with monocyte-enriched proteins with correlation larger than 0.8 or less than -0.8.**

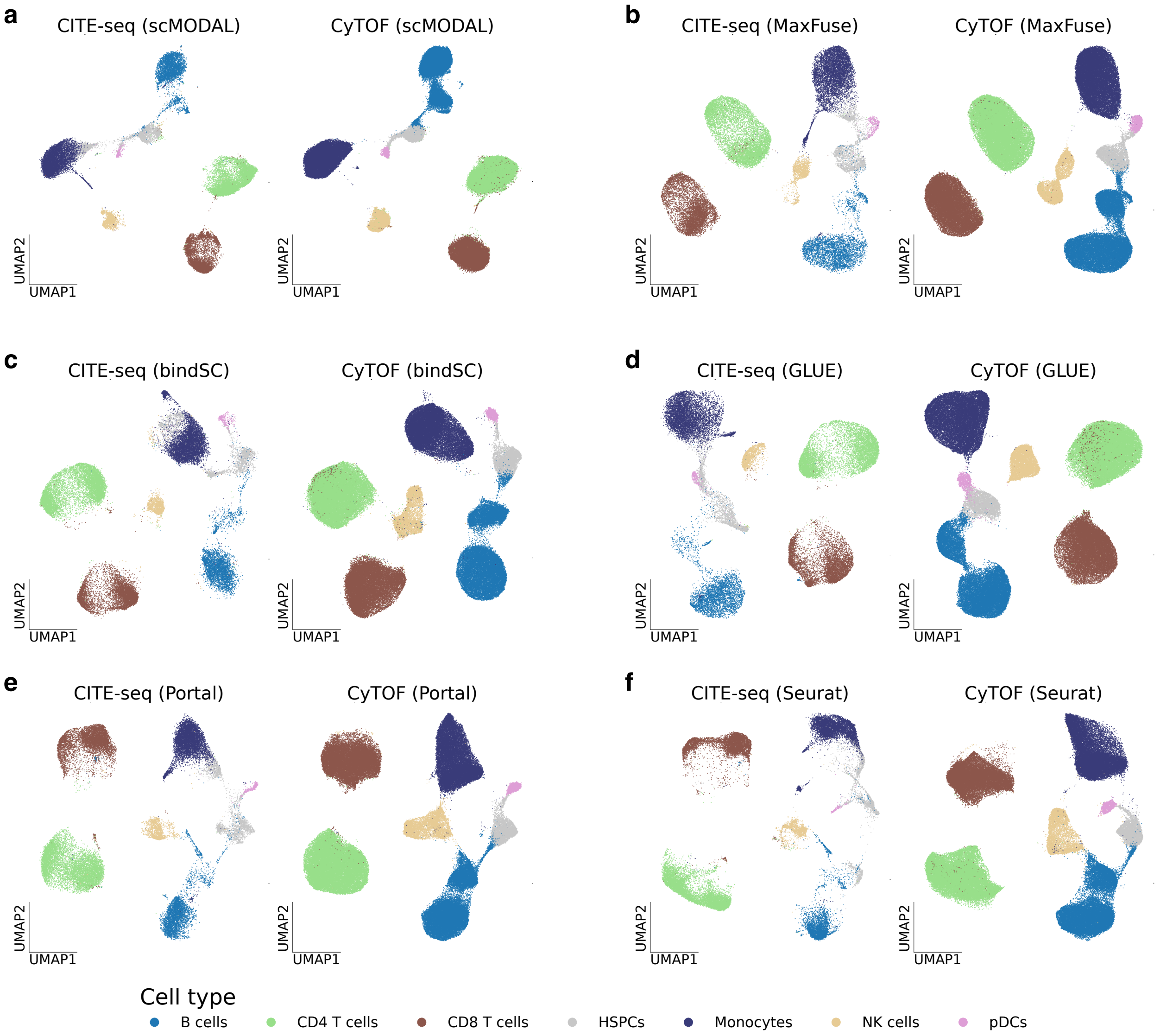


Supplementary Fig. 4: **Comparison of integration methods based on the CITE-seq [8] and the CyTOF [9] human bone marrow data. a-f**. UMAP plots of integrated embeddings produced by scMODAL (**a**), MaxFuse (**b**), bindSC (**c**), GLUE (**d**), Portal (**e**) and Seurat (**f**), colored by cell types.

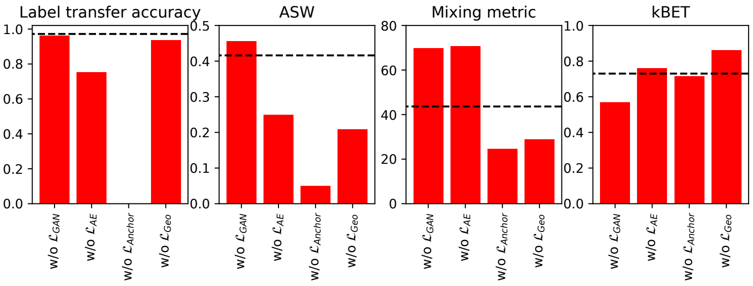


Supplementary Fig. 5: **Evaluation of scMODAL's performance by removing its components, using the CITE-seq and the CyTOF human bone marrow data.** Horizontal dashed lines indicate the original model performance. For all metrics except the mixing metric, a higher score indicates a better performance. Notabily, without the MNN anchor regularization, the label transfer accuracy is very close to 0 because the cell types are wrongly matched.

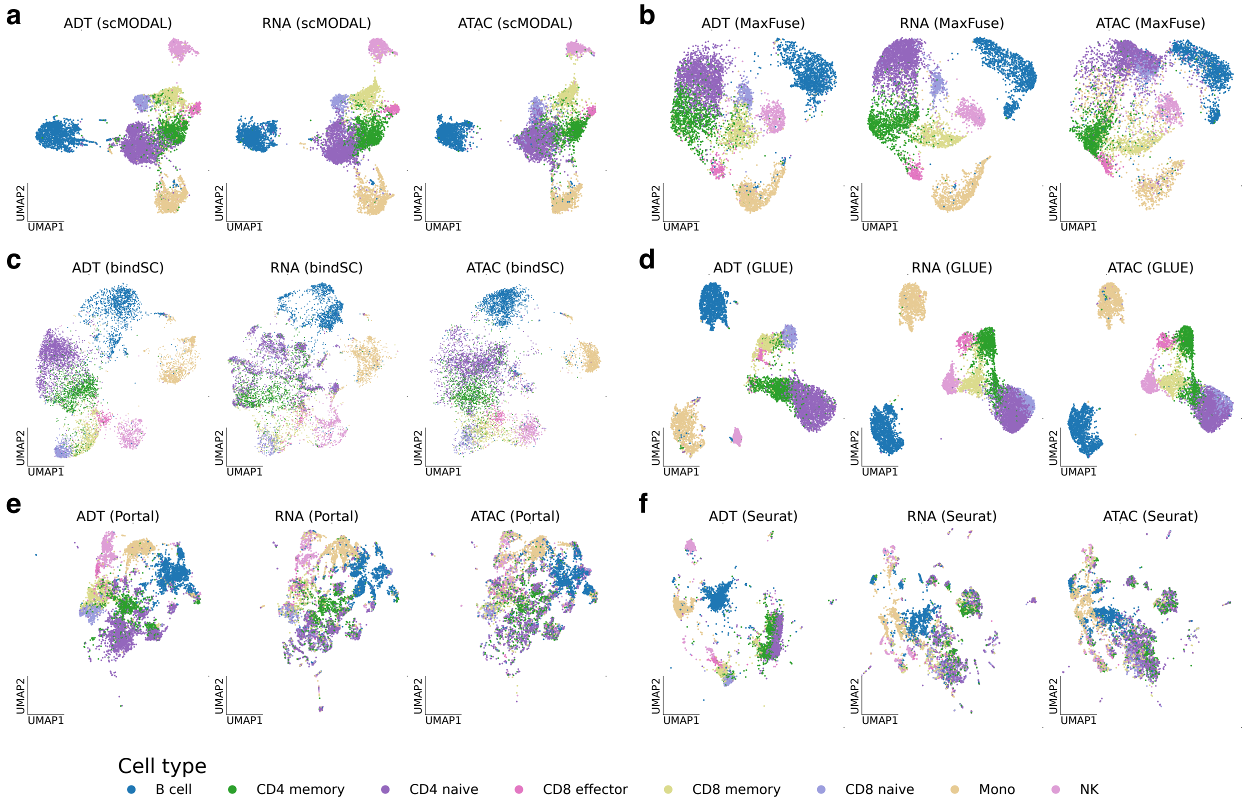


Supplementary Fig. 6: **Comparison of integration methods based on the TEA-seq human PBMC data [10]. a-f**. UMAP plots of integrated embeddings produced by scMODAL (**a**), MaxFuse (**b**), bindSC (**c**), GLUE (**d**), Portal (**e**) and Seurat (**f**), colored by cell types.

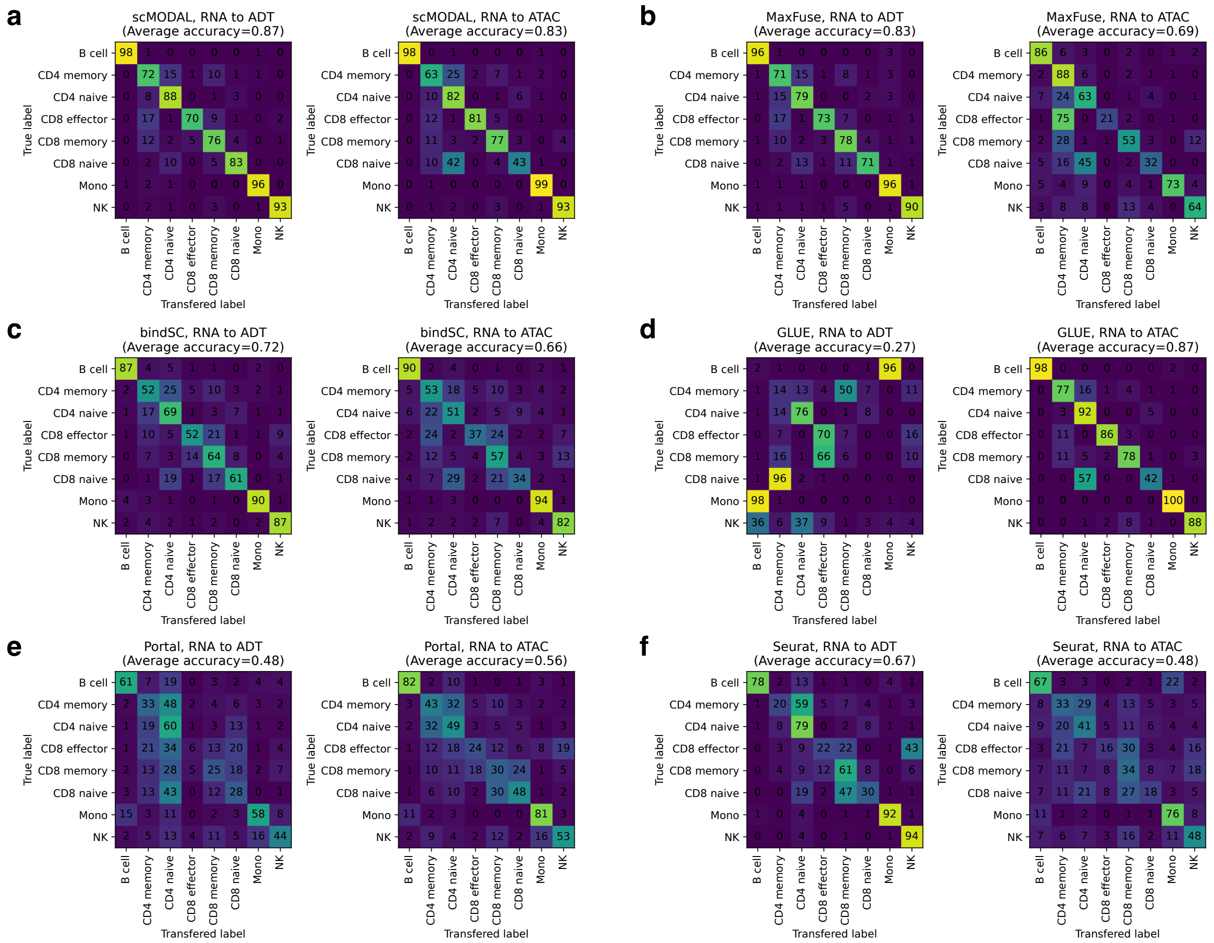


Supplementary Fig. 7: **Confusion matrices of the label transfer results based on the TEA-seq human PBMC data. a-f.** Confusion matrices produced by scMODAL (**a**), MaxFuse (**b**), bindSC (**c**), GLUE (**d**), Portal (**e**) and Seurat (**f**).

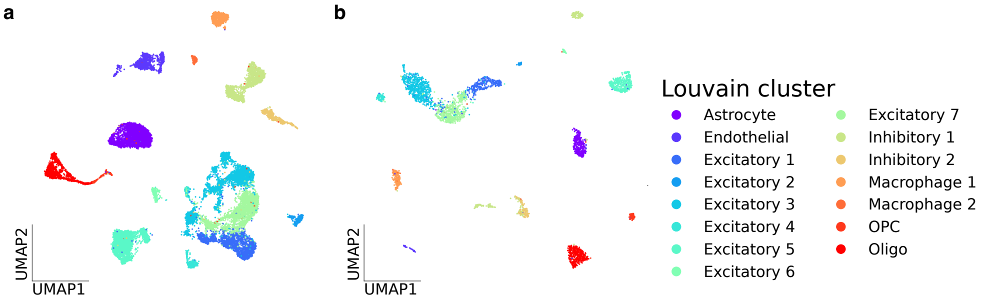


Supplementary Fig. 8: **UMAP plots of unintegrated scRNA-seq (a) [11] and scATAC-seq (b) mouse brain datasets, colored by Louvain [12] cluster labels.**

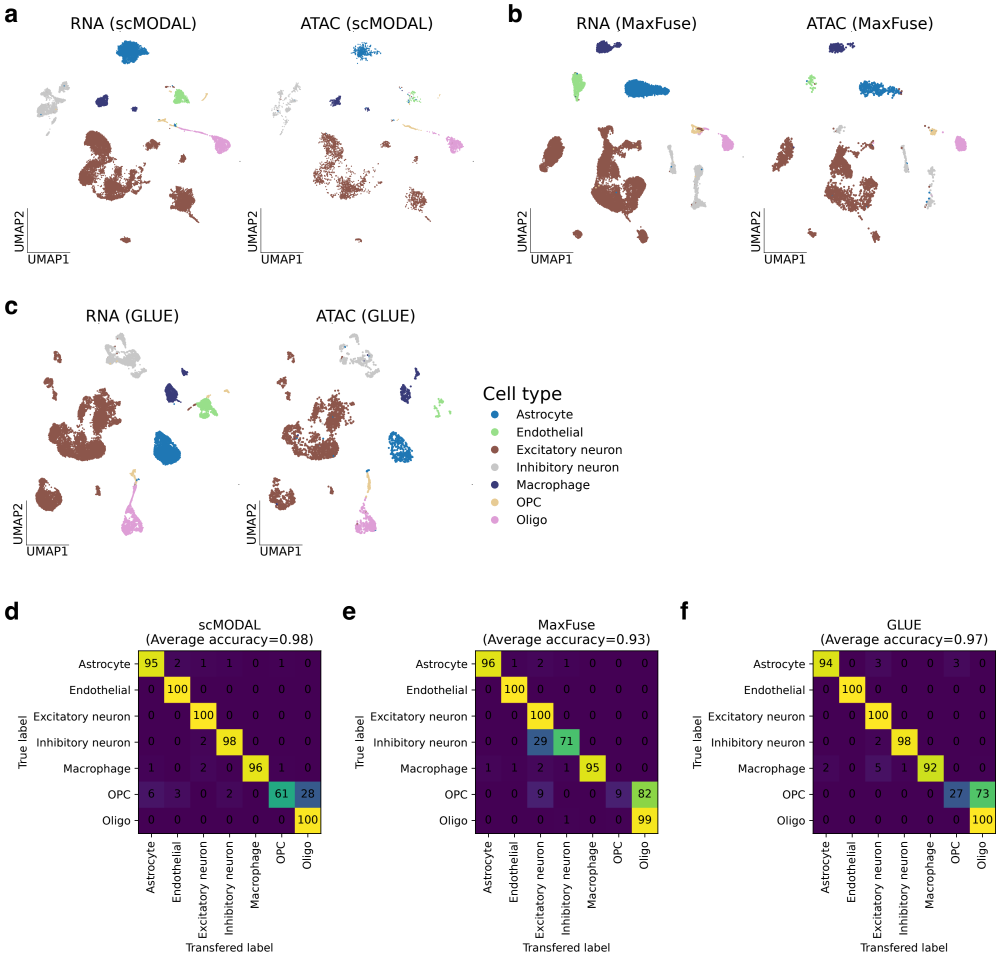


Supplementary Fig. 9: **Integration of the scRNA-seq and scATAC-seq mouse brain datasets. a-c**. UMAP plots of integrated embeddings produced by scMODAL (**a**), MaxFuse (**b**) and GLUE (**c**), colored by cell types. **d-f**. Confusion matrices produced by scMODAL (**d**), MaxFuse (**e**) and GLUE (**f**).

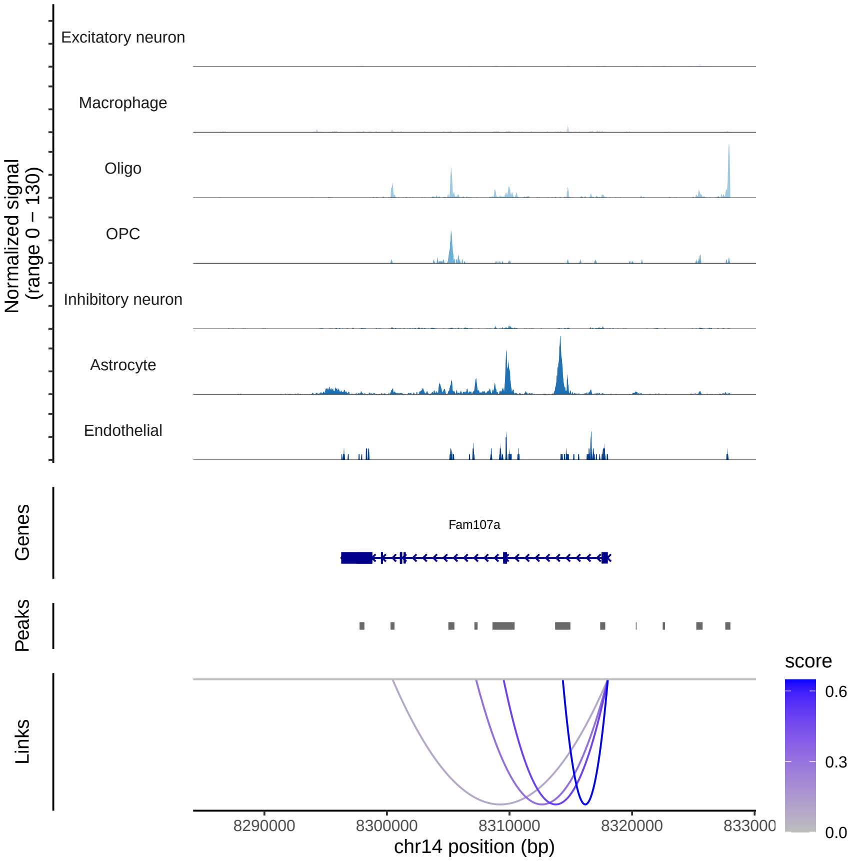


Supplementary Fig. 10: **Visualization of the mouse brain scATAC-seq data at the *Fam107a* locus.** Top panel: ATAC-seq tracks at the *Fam107a* locus. Bottom panel: gene-peak links inferred by Signac [13].

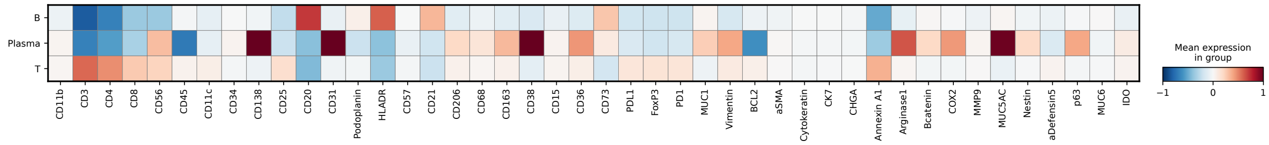


Supplementary Fig. 11: **Abundance levels of measured proteins in the human tonsil CODEX dataset [14].**

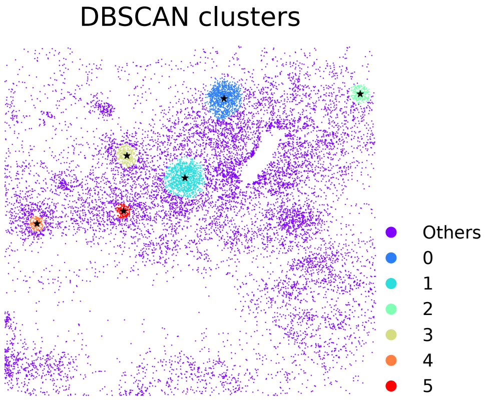


Supplementary Fig. 12: **Six DBSCAN [15] clusters identified using the spatial distribution of B-Ki67 cells.**

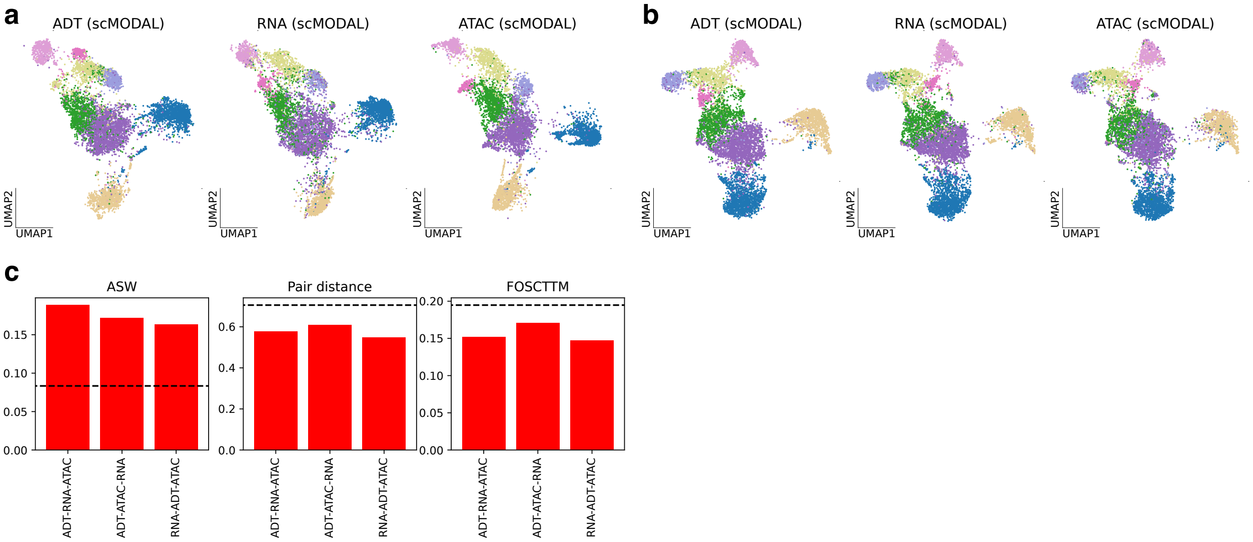


Supplementary Fig. 13: **Comparison of different integration orders using the TEA-seq human PBMC data. a**, **b**. UMAP plots of integrated embeddings produced by scMODAL using the integration orders ADT-ATAC-RNA (**a**) and RNA-ADT-ATAC (**b**), colored by cell types. The original integration order was set to be ADT-RNA-ATAC. **c**. ASW, pair distance, and FOSCTTM metrics are shown for all integration orders. Horizontal dashed lines represent the second-best method, MaxFuse. For ASW, higher values indicate better performance, while for pair distance and FOSCTTM, lower values are preferable.

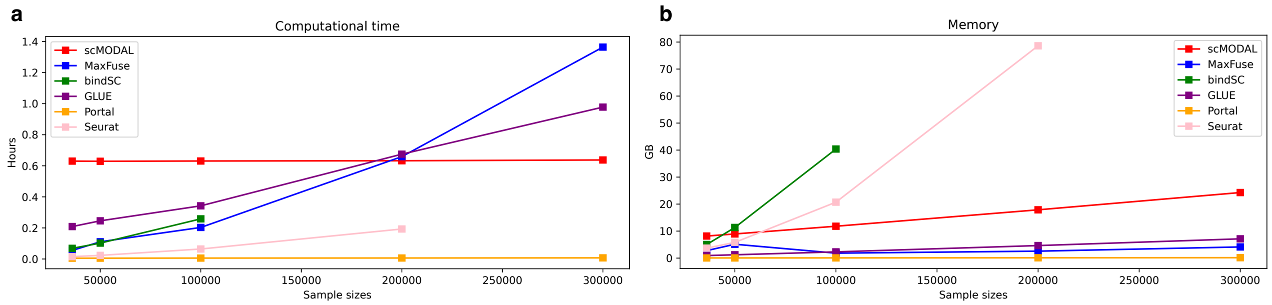


Supplementary Fig. 14: **Computational time and peak memory usage.** We used the CITE-seq human PBMC data to compare the computational time (**a**) and peak memory usage (**b**) of all compared methods with varying sample sizes.

### Supplementary Tables

#### Network structures in scMODAL

| Layer | Detail | Input Size | Output Size | Number of Parameters |
|---|---|---|---|---|
| Fully Connected | Linear | $n$ | 512 | $512n + 512 = 512(n + 1)$ |
| | RELU | 512 | 512 | 0 |
| Fully Connected | Linear | 512 | 20 | $512 \times 20 + 20 = 10,260$ |

Supplementary Table 1: **Structure of scMODAL's encoder networks.** The number of parameters depends on the number of input features $n$.

| Layer | Detail | Input Size | Output Size | Number of Parameters |
|---|---|---|---|---|
| Fully Connected | Linear | 20 | 512 | $20 \times 512 + 512 = 10,752$ |
| | RELU | 512 | 512 | 0 |
| Fully Connected | Linear | 512 | 30 | $512n + n = 513n$ |

Supplementary Table 2: **Structure of scMODAL's generator networks.** The number of parameters depends on the number of input features $n$.

| Layer | Detail | Input Size | Output Size | Number of Parameters |
|---|---|---|---|---|
| Fully Connected | Linear | 20 | 512 | $20 \times 512 + 512 = 10,752$ |
| | RELU | 512 | 512 | 0 |
| Fully Connected | Linear | 512 | 512 | $512 \times 512 + 512 = 262,656$ |
| | RELU | 512 | 512 | 0 |
| Fully Connected | Linear | 512 | 1 | $512 \times 1 + 1 = 513$ |

Supplementary Table 3: **Structure of scMODAL's discriminator networks.**

### References


[1] Hao, Y. *et al.* Integrated analysis of multimodal single-cell data. *Cell* **184**, 3573–3587.e29 (2021).

[2] McInnes, L., Healy, J., Saul, N. & Grossberger, L. UMAP: Uniform manifold approximation and projection. *The Journal of Open Source Software* **3**, 861 (2018).

[3] Chen, S. *et al.* Integration of spatial and single-cell data across modalities with weakly linked features. *Nature Biotechnology* **42**, 1096–1106 (2024).

[4] Dou, J. *et al.* Bi-order multimodal integration of single-cell data. *Genome Biology* **23**, 112 (2022).

[5] Cao, Z.-J. & Gao, G. Multi-omics single-cell data integration and regulatory inference with graph-linked embedding. *Nature Biotechnology* **40**, 1458–1466 (2022).

[6] Zhao, J. *et al.* Adversarial domain translation networks for integrating large-scale atlas-level single-cell datasets. *Nature Computational Science* **2**, 317–330 (2022).

[7] Stuart, T. *et al.* Comprehensive integration of single-cell data. *Cell* **177**, 1888–1902.e21 (2019).

[8] Stuart, T. & Satija, R. Integrative single-cell analysis. *Nature Reviews Genetics* **20**, 257–272 (2019).

[9] Levine, J. H. *et al.* Data-driven phenotypic dissection of AML reveals progenitor-like cells that correlate with prognosis. *Cell* **162**, 184–197 (2015).

[10] Swanson, E. *et al.* Simultaneous trimodal single-cell measurement of transcripts, epitopes, and chromatin accessibility using TEA-seq. *Elife* **10**, e63632 (2021).

[11] Zeisel, A. *et al.* Molecular architecture of the mouse nervous system. *Cell* **174**, 999–1014.e22 (2018).

[12] Blondel, V. D., Guillaume, J.-L., Lambiotte, R. & Lefebvre, E. Fast unfolding of communities in large networks. *Journal of Statistical Mechanics: Theory and Experiment* **2008**, P10008 (2008).

[13] Stuart, T., Srivastava, A., Madad, S., Lareau, C. A. & Satija, R. Single-cell chromatin state analysis with Signac. *Nature Methods* **18**, 1333–1341 (2021).

[14] Brbić, M. *et al.* Annotation of spatially resolved single-cell data with STELLAR. *Nature Methods* **19**, 1411–1418 (2022).

[15] Ester, M., Kriegel, H.-P., Sander, J., Xu, X. *et al.* A density-based algorithm for discovering clusters in large spatial databases with noise. In *kdd*, vol. 96, 226–231 (1996).
